# Lateral hypothalamic GABAergic neurons encode alcohol memories

**DOI:** 10.1101/2023.11.16.567157

**Authors:** Isis Alonso-Lozares, Pelle Wilbers, Lina Asperl, Sem Teijsse, Charlotte van der Neut, Dustin Schetters, Yvar van Mourik, Allison J. McDonald, Tim Heistek, Huibert D. Mansvelder, Taco J. De Vries, Nathan J. Marchant

## Abstract

In alcohol use disorder the alcohol memories persist during abstinence, and exposure to stimuli associated with alcohol use can lead to relapse. This highlights the importance of investigating the neural substrates underlying not only relapse, but also encoding and expression of alcohol memories. GABAergic neurons in the Lateral Hypothalamus (LH-GABA) have been shown to be critical for food-cue memories and motivation, however the extent to which this role extends to alcohol-cue memories and motivations remains unexplored. In this study, we aimed to describe how alcohol related memories are encoded and expressed in LH GABAergic neurons. Our first step was to monitor LH-GABA calcium transients during acquisition, extinction, and reinstatement of an alcohol-cue memory using fiber photometry. We trained the rats on a Pavlovian conditioning task where one conditioned stimulus (CS+) predicted alcohol (20% EtOH) and another conditioned stimulus (CS-) had no outcome. We then extinguished this association through non-reinforced presentations of the CS+ and CS-, finally in two different groups we measured relapse under non-primed and alcohol primed induced reinstatement. Our results show that initially both cues caused increased LH-GABA activity, and after learning only the alcohol-cue increased LH-GABA activity. After extinction this activity decreases, and we found no differences in LH-GABA activity during reinstatement in either group. Next, inhibited LH-GABA neurons with optogenetics to show that activity of these neurons is necessary for the formation of an alcohol-cue associations. These findings suggest that LH-GABA might be involved in attentional processes modulated by learning.

## Introduction

Alcohol is one of the most widely available and frequently consumed substances in our society^1^. While most individuals (80-90%) can consume alcohol in a moderate and controlled manner, a small percentage develop alcohol use disorder (AUD). In the European Union alone, this disorder affects approximately 23 million people^2^. AUD is characterized by persistent alcohol use despite negative consequences and a high likelihood of relapse^3^.

Individuals with AUD often go through multiple cycles of treatment, periods of abstinence, and relapse before achieving full recovery^4^.

Relapse can occur when an individual encounters cues and environments associated with previous alcohol use^5,6^, and these experiences can cause craving^7^ contributing to relapse^8^. These strong positive associations between neutral stimuli (e.g., a bar, a specific location, certain individuals) and alcohol are formed in the initial stages of addiction development and share many similarities with behavioural patterns triggered by natural rewards like food^9^. Like natural rewards, drugs create anticipation of positive outcomes and reinforce the individual’s motivation to seek them. Consequently, pathological associative learning is a critical component of the addiction cycle, where the brain’s mechanisms that typically promote survival-related behaviours are hijacked by a drug’s more subjectively potent stimuli^10^. It is therefore crucial to better understand how these memories are formed in the brain.

The Lateral Hypothalamus (LH) has been considered a critical substrate of appetitive and consummatory behaviours since it was discovered that lesions cause hypophagia, while stimulation has the opposite effect^11,12^. Others pointed at the LH having a general role in motivation^13,14^, and more recent data suggests the role of LH extends beyond promoting feeding behaviour to orchestrating complex learning processes related to the encoding and maintenance of motivationally relevant cue-food associations^15,16,17^, as well as other learning processes^18,19^. In relation to relapse, increased activity in LH is associated with context-induced reinstatement of alcohol seeking after both extinction and punishment^20,21^, as well as context-induced reinstatement of extinguished cocaine seeking^22^.

Recent evidence suggests that the GABAergic subpopulation of LH (LH-GABA) are involved in associative learning processes and motivated behaviours. These neurons are functionally and anatomically distinct from the Orexin (Ox) and Melanin-Concentrating Hormone (MCH) populations^16,23^, and have reciprocal projections with Ventral Tegmental Area^24^ (VTA) and Nucleus Accumbens Shell^25^ (NAcS), areas well known for their roles in reward learning. LH-GABA activity is associated with appetitive or consummatory behaviors^16^, and opto-inhibition of LH-GABA subpopulation prevents learning of cue-food associations^17^.

Currently the role of LH-GABA in alcohol related learning is unknown. Here, we trained rats on an alcohol Pavlovian conditioning task and used fiber photometry calcium imaging^26,27^ to show that LH-GABA activity is initially high to cues when they are delivered, and across learning this activity is sustained to the alcohol-predictive cue but decreases to the non-predictive cue. We also show in extinction that activity in LH-GABA neurons track violations in expected outcomes, with sustained activity when alcohol is expected but not delivered. To investigate a causal role of LH-GABA neurons we used optogenetics^28^ to inhibit activity during conditioning and found that acquisition is decreased during cue presentation. These data highlight the importance of LH-GABA in both encoding and expression of alcohol associated memories.

## Results

### Exp. 1: Validation of the dual-virus approach to target GABAergic neurons in LH

To target the GABAergic subpopulation of LH neurons we used dual-virus approach involving combination of a GAD-Cre virus with different Cre-dependent viruses^29^. To first validate this approach, we injected a cocktail of one AAV encoding Cre driven by the GAD promoter with another AAV encoding Cre-dependent eGFP. Immunofluorescence labelling MCH and OX-expressing neurons revealed minimal overlap between both OX- immunoreactive (Figure 1B) and MCH-immunoreactive (Figure 1C) neurons and GFP expression driven under the GAD promoter. Using RNA-scope to label GAD mRNA (Figure 1A), we found high overlap between GAD+ and eGFP (Figure 1D). These data show that the dual-virus approach was successful in confining expression of AAV-induced transgenes to GABAergic neurons.

**Figure 1.**
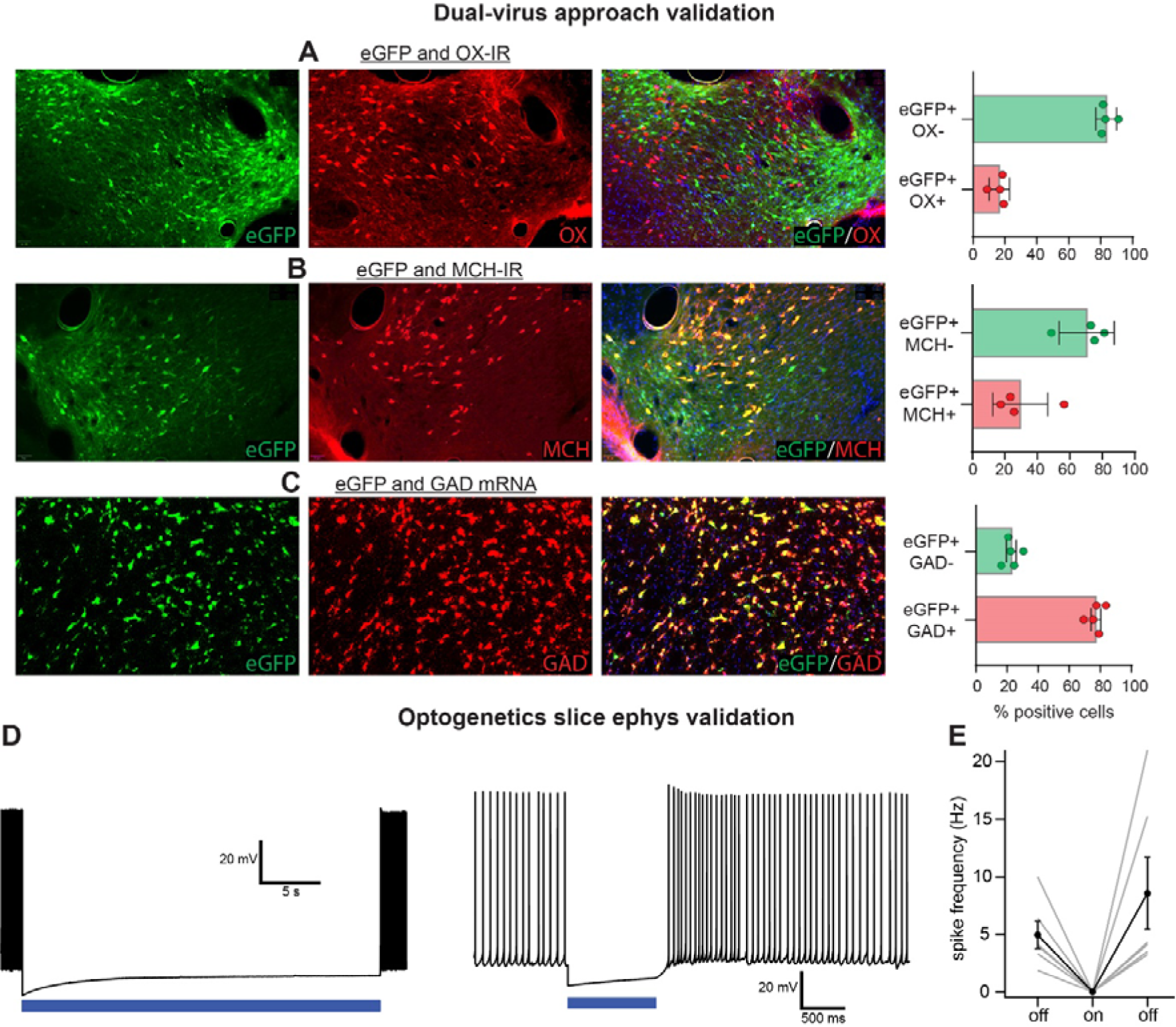
Validation of the dual-virus approach to target LH-GABA neurons, and that blue light inhibits GtACR2 expressing LH-GABA neurons. Representative images of eGFP expression and co-labelling of OX (A), MCH (B) and GAD (C). Data are mean (+/- SEM; n=4 for MCH and OX validation, n=5 for GAD validation) % of eGFP+ only or eGFP+OX (A), eGFP+MCH (B), eGFP+GAD (C). (D) Example of light-evoked suppression of neuronal firing inGtACR2 expressing neurons. (E) Summary data plotting mean (+/- SEM; n=6) spike frequency in the presence and absence of light stimulation. Blue bars indicate light delivery.

To validate that blue light effectively inhibits the inhibitory opsin GtACR2^28^ expressing neurons, we made whole cell recordings from fluorescent labelled neurons (Figure 1D-E). Current injection into GtACR2 expression cells depolarized them above spike threshold, and activating GtACR2 with blue light significantly decreased the rate of current-induced action potentials (n=6).

### Exp. 2: Monitoring LH-GABA activity during alcohol conditioning, extinction, and primed reinstatement of alcohol-seeking

In experiment 2, we aimed to identify LH-GABA activity during alcohol conditioning, extinction, and primed reinstatement of alcohol seeking. Figure 2A shows the experimental timeline, the expression of the Ca2+ sensor jGCaMP7f^30^ in LH-GABA was achieved using a mixture of one AAV encoding Cre driven by the GAD promoter with another AAV encoding Cre-dependent jGCaMP7f under the hSyn1 promoter. We measured calcium (Ca2+) transients via an optic fiber cannula implanted above LH throughout the entire task (Figure 2B). Experiment 2 is comprised of two cohorts of rats. Cohort 1 (n= 2 male rats, n=3 female rats; Exp. 2a) received the targeting and behavioral strategies described above. Cohort 2 (n=13 female rats; Exp. 2b) received the targeting and behavioral strategies described above as well as additional chemogenetic viral infections, and additional test sessions with a chemogenetic ligand deschloroclozapine. The outcome of the chemogenetic tests were negative and therefore we combined the analysis for these two cohorts because the methodology was otherwise identical. Specific details of the effects of the chemogenetic tests on alcohol seeking and LH-GABA activity are described in the Supplementary Materials.

**Figure 2.**
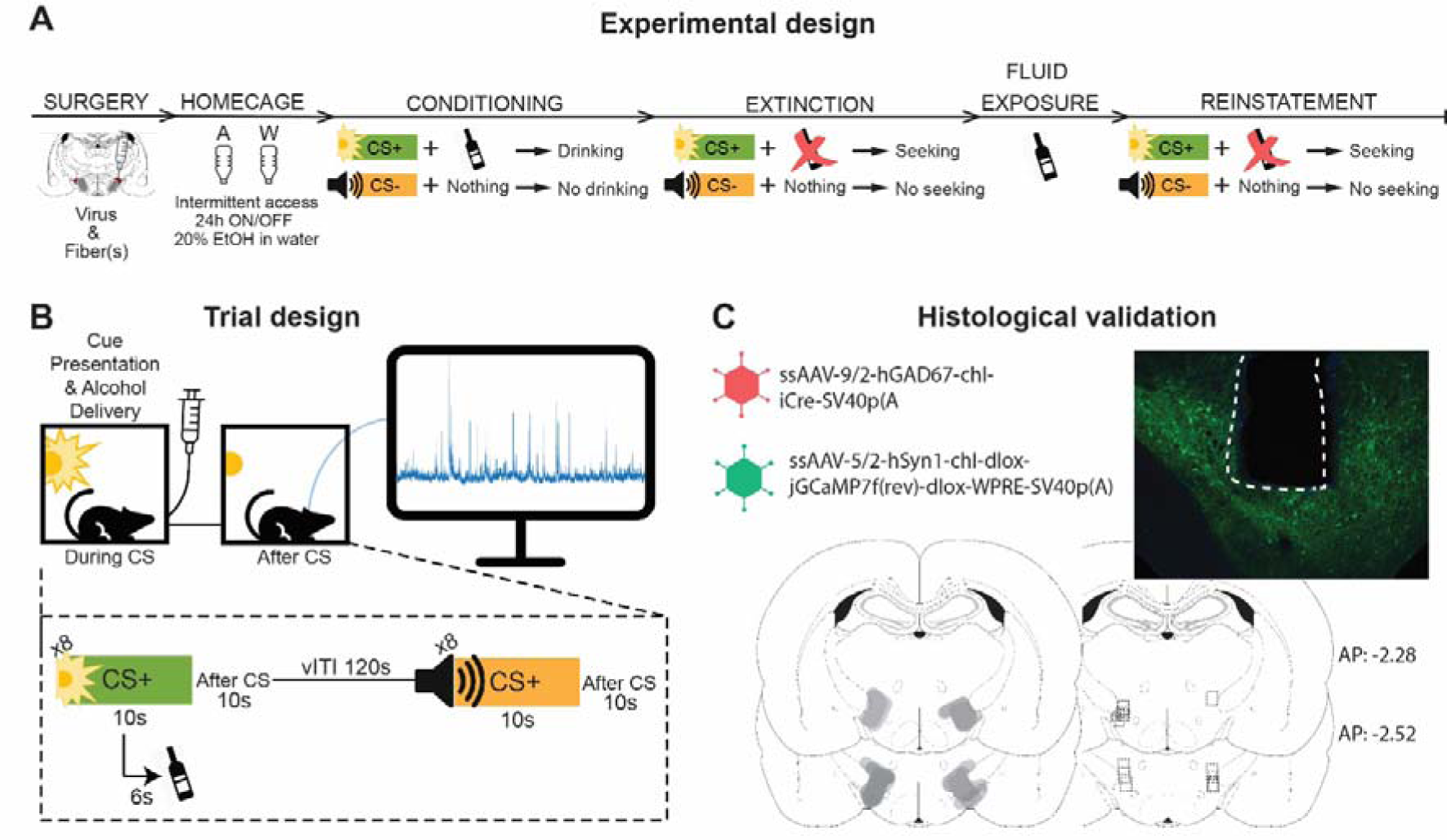
Experimental design and histological validation of fiber implants and jGCaMP7f expression in LH. (A) Outline of the experimental procedure (n =18; 2 males, 16 females). (B) Trial design for photometry experiments. (C) Representative image of jGCaMP7f expression and fiber implant in LH, and fiber placement and expression validation.

#### Across learning, LH-GABA neuron activity increases to an alcohol-predictive cue and decreases to a non-relevant cue

<u>Behavioural data:</u> We used repeated measures ANOVA with the within-subjects factors Session (First, Last) and CS Type (CS+, CS-) to compare alcohol seeking during the first and the last conditioning sessions (n=19 sessions; Figure 3A and 3B). We found main effects of Session and CS Type, as well as a significant Session x CS Type interaction on time spent in the magazine both during (Figure 3B; F(1, 17)=133,7, p < 0.0001) and after (Figure 3C; F(1, 17)=4747, p < 0.0001) CS presentation. We found no difference between CS Type on time in the magazine during pre-CS (S2). These data show that the rats learned to discriminate between the alcohol-predictive and neutral cues, and alcohol seeking increased during the CS+ but not the CS-.

**Figure 3.**
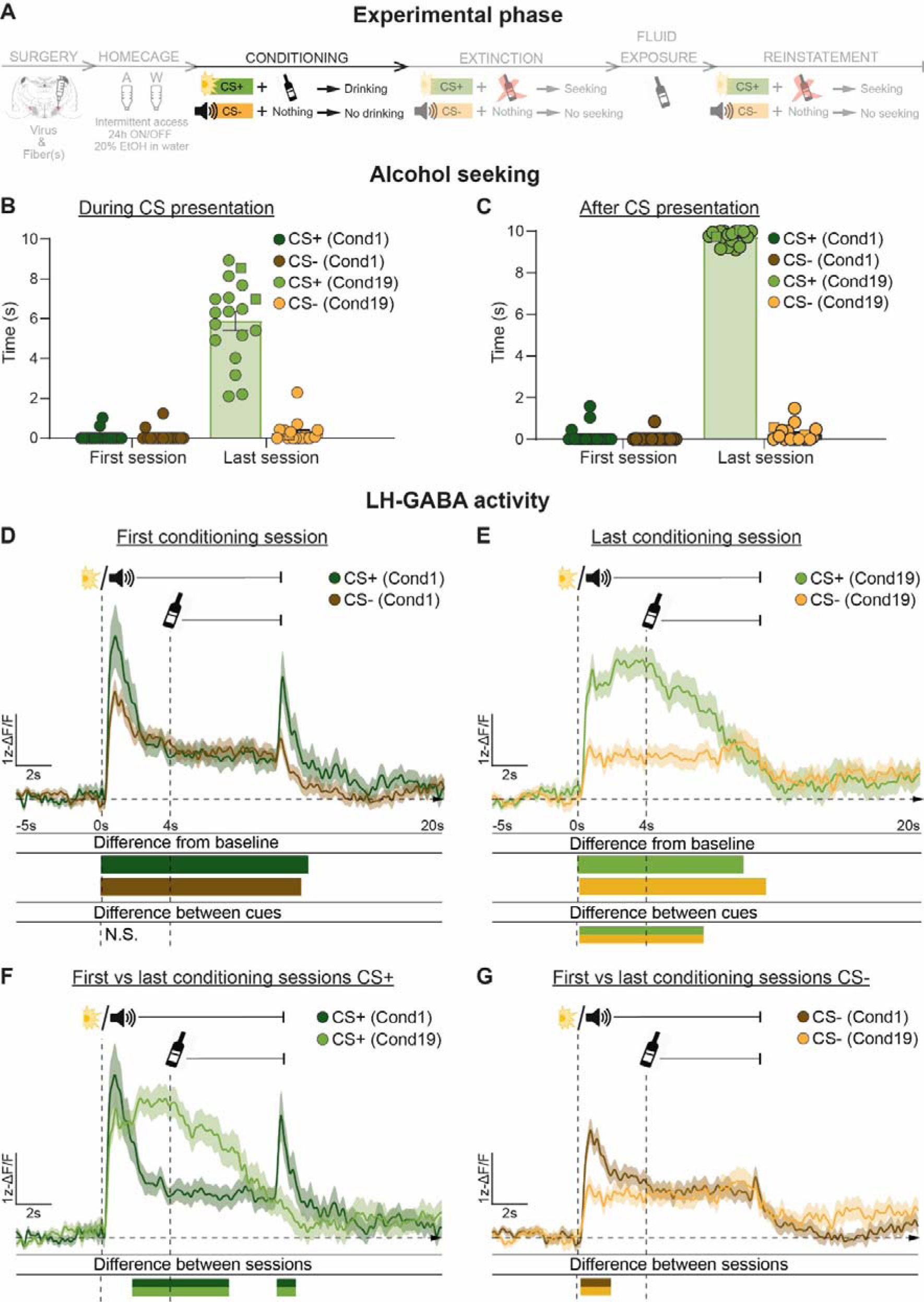
Monitoring LH-GABA during alcohol Pavlovian conditioning. (**A**) Outline of experimental procedure (n=2 males, n=16 females). Mean ± standard error of the mean (SEM) time spent in the alcohol magazine during **(B)** and after **(C)** CS presentation comparing the first and last conditioning sessions (n=8 trials per CS type, per session, individual data points show males (squares) and females (dots)). Alcohol was delivered at 4 s after CS onset. Ca2+ traces of LH-GABA activity centered around cue onset (-5s to +20s) comparing LH-GABA activity to CS+ and CS- on the first **(D)** and last **(E)** conditioning sessions. (CS+ Cond1: n=169; CS- Cond1: n=168; CS+ Cond19: n =170; CS- Cond19: n=169). The same mean Ca2+ traces are plotted for comparisons across sessions between CS+ **(F)** and CS-**(G)**. For all photometry traces, bars at bottom of graph indicate significant deviations from baseline (dF/F ≠ 0) determined via bootstrapped confidence intervals (99% CIs), or significant differences between the specific events (CS+ Cond1; CS- Cond1; CS+ Cond19; CS- Cond19) determined via permutation tests with alpha 0.01 for comparisons between CS type and session. Vertical dashed lines indicate CS onset and alcohol delivery for CS+, horizontal line indicates baseline (dF/F=0).

<u>Photometry data:</u> We made comparisons within a single session between the CS types (CS+ vs CS-), and across the different sessions within the same CS type (First vs. Last; Figure 3. D-G). Detailed time-points for specific significance time windows can be found on Supplementary Table 1. In the first conditioning session (Figure 3D), we found that LH-GABA activity was significantly higher for both CS types, with no difference between the two. In the last conditioning session (Figure 3E), we observed a significant increase from baseline for CS+ and CS-. In contrast to the first session, we found activity to the CS+ was significantly higher than to the CS-. These data suggest that acquisition of a cue-alcohol memory is associated with increased activity of LH-GABA neurons. To further investigate this, we compared activity to the CS+ or CS- between the first and last conditioning sessions. We found that acquisition of the cue-alcohol memory was associated with both longer sustained activity to the CS+ (Figure 3F), and decreased activity to the CS- (Figure 3G). Overall, these data show that the activity of LH-GABA neurons is bidirectionally shaped by learning experience. We found both an increase in activity to a motivationally relevant (alcohol-predictive) cue, as well as decreased activity to a motivationally non-relevant cue (CS-).

#### Expression of the alcohol memory

<u>Behavioural data:</u> Repeated measures ANOVA comparing the Last conditioning and First Extinction sessions (Figure 4B and 4C) revealed a significant Session x CS Type interaction on time spent in the magazine both during (Figure 4B;F(1, 17)=166,0, p < 0.0001) and after (Figure 4C; F(1, 17)=1772, p < 0.0001) CS presentation. Thus, we found that rats spent significantly more time in the alcohol port during CS+ than CS- in the absence of alcohol delivery, suggesting the learned memory was expressed without the alcohol outcome.

**Figure 4.**
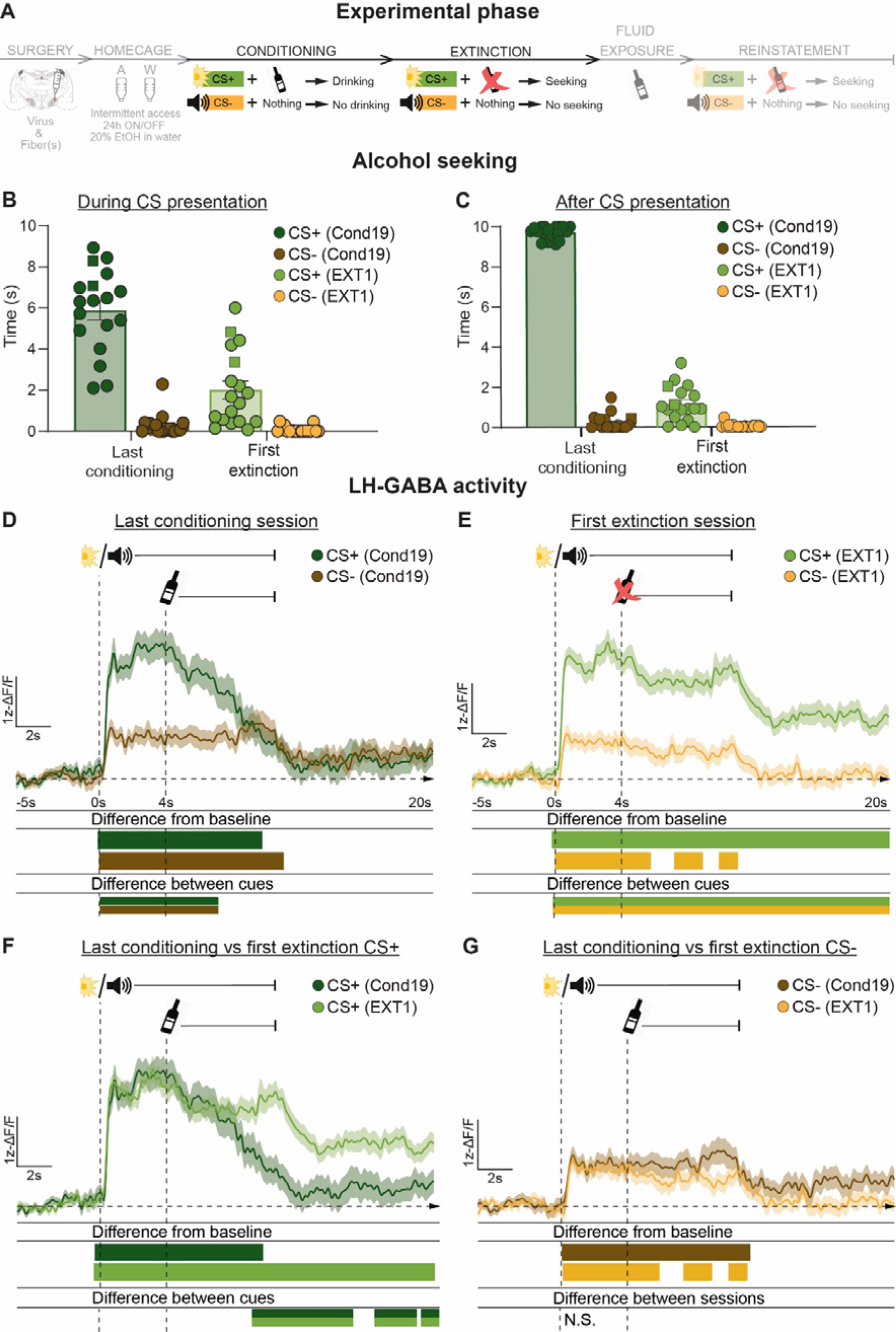
Monitoring LH-GABA during alcohol Pavlovian conditioning and extinction. (**A**) Outline of experimental procedure (n=2 males, n=16 females). Mean ± standard error of the mean (SEM) time spent in the alcohol magazine during (**B**) and after (**C**) CS presentation comparing the last conditioning and first extinction sessions (n=8 trials per CS type, per session, individual data points show males (squares) and females (dots)). Alcohol was delivered at 4 s after CS onset. Ca2+ traces of LH-GABA activity centred around cue onset (-5s to +20s) comparing LH-GABA activity to CS+ and CS- on the last conditioning (**D**) and first extinction (**E**) sessions. (CS+ Cond19: n=170; CS- Cond19: n=169; CS+ EXT1: n=61; CS- EXT1: n=136). The same mean Ca2+ traces are plotted for comparisons across sessions between CS+ (**F**) and CS-(**G**). For all photometry traces, bars at bottom of graph indicate significant deviations from baseline (dF/F ≠ 0), or significant differences between the specific events (CS+ Cond19; CS- Cond19; CS+ EXT1; CS- EXT1), determined via bootstrapped confidence intervals (99% CIs), and permutation tests with alpha 0.01 for comparisons between CS type and session. Vertical dashed lines indicate CS onset and alcohol delivery for CS+, horizontal line indicates baseline (dF/F=0).

<u>Photometry data:</u> We made comparisons within a single session between CS types (CS+ vs CS-), and across the different sessions within the same CS type (First vs. Last; Figure 4. D-G). Here we plotted the same data from Figure 3D into Figure 4D, and results for the last conditioning session are described above. Detailed time-points for specific significance time windows can be found on Supplementary Table 1. On the first extinction session (Figure 4E), we observed a significant increase from baseline for both cues. Comparison of activity between CS types revealed that activity to CS+ remains significantly higher compared to CS- both during and after cue presentation. To determine if LH-GABA activity decreases during extinction we compared activity during the last conditioning and first extinction sessions for CS+ (Figure 4F) or CS- (Figure 4G). Interestingly, while we found no difference in CS+ or CS- induced activity when the cues were delivered, there was higher activity after the CS+ in the first extinction session compared to the last conditioning session. This sustained activity was during the period when alcohol is expected but not delivered.

#### Extinction

<u>Behavioural data:</u> Repeated measures ANOVA comparing alcohol seeking across trials on the First Extinction session (n=8 trials per CS; Fig. S6B and Fig. S6C), revealed a significant Trial x Type of CS interaction both during (F(7, 119)=3,694, p < 0.01), and after (F(7, 119)=4,239, p < 0.001). This suggests extinction of seeking is observed already within the first session. Repeated measures ANOVA comparing the first and the last extinction sessions (n=7 sessions; Figure 5B and 5C) revealed a significant Session x CS Type interaction on time spent in the magazine both during (Figure 5B; F(1, 17)=19,96, p < 0.001) and after (Figure 5C; F(1, 17)=34,75, p < 0.0001) CS presentation. Thus, we show that rats spent significantly more time in the alcohol port during and after CS+ onset on the first extinction session compared to CS-. This contrasts with the last extinction session, where we show that time spent in the alcohol port did not significantly differ between CS+ and CS-.

**Figure 5.**
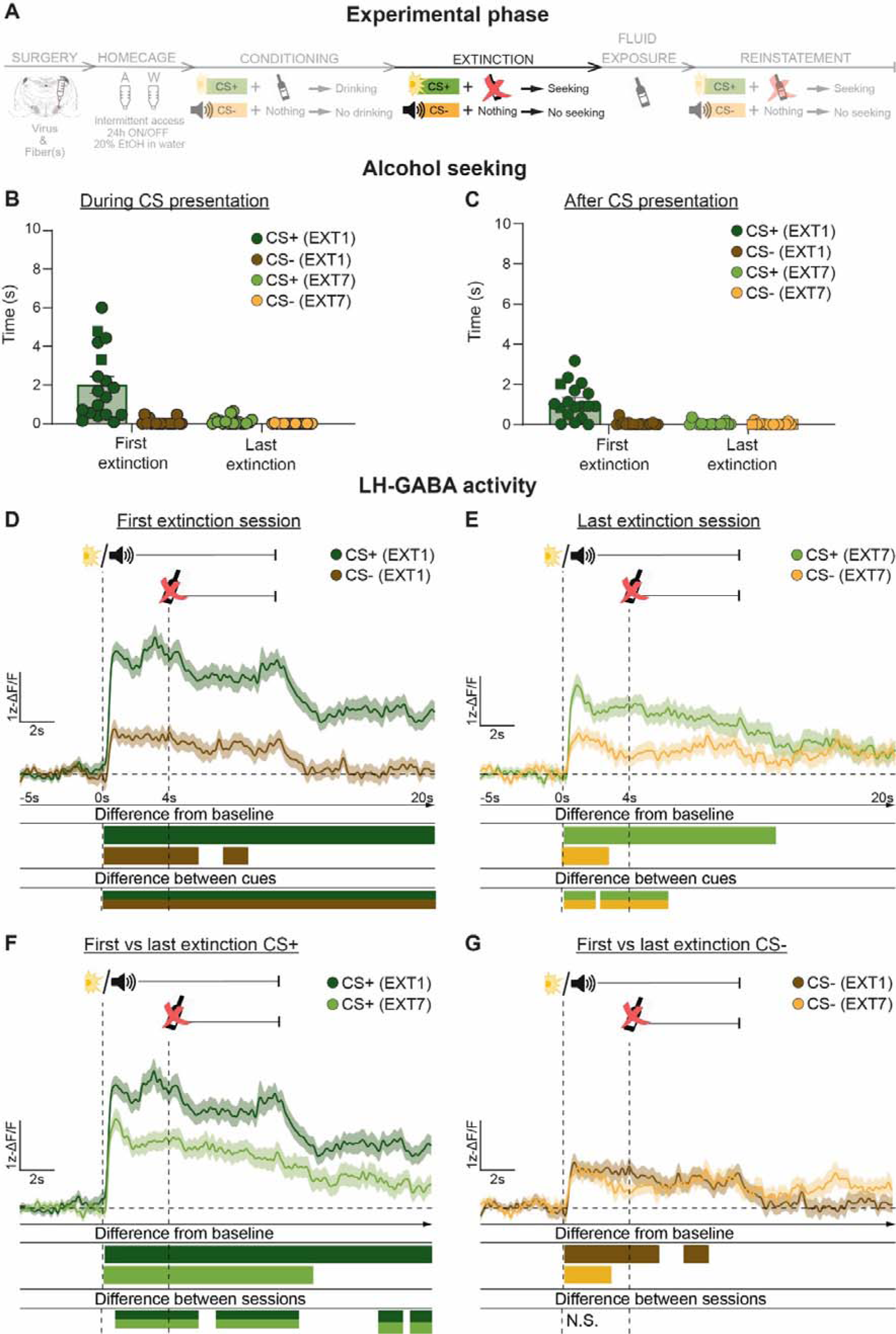
Monitoring LH-GABA during extinction. (**A**) Outline of experimental procedure (n=2 males, n=16 females). Mean ± standard error of the mean (SEM) time spent in the alcohol magazine during (**B**) and after (**C**) CS presentation comparing the first and last extinction sessions. (n=8 trials per CS type, per session, individual data points show males (squares) and females (dots)). Ca2+ traces of LH-GABA activity centred around cue onset (-5s to +20s) comparing LH-GABA activity to CS+ and CS- on the first (**D**) and last (**E**) extinction sessions. (CS+ EXT1: n=161; CS- EXT1: n=136; CS+ EXT7: n=161; CS- EXT7: n=136). The same mean Ca2+ traces are plotted for comparisons across sessions between CS+ (**F**) and CS- (**G**). For all photometry traces, bars at bottom of graph indicate significant deviations from baseline (dF/F ≠ 0), or significant differences between the specific events (CS+ EXT1; CS- EXT1; CS+ EXT7; CS- EXT7), determined via bootstrapped confidence intervals (99% CIs), and permutation tests with alpha 0.01 for comparisons between CS type and session. Vertical dashed lines indicate CS onset and alcohol delivery for CS+, horizontal line indicates baseline (dF/F=0).

<u>Photometry data:</u> We made comparisons between the first 4 and last 4 trials of the First Extinction session for CS+ and CS- (Fig. S6D-G) and found no significant differences in LH-GABA activity. From this we interpret that learning changes in LH cannot be identified within a single session. Therefore, we made comparisons within a single session between the different CS types (CS+ vs CS-), and across the different sessions within the same CS type (First vs. Last; Figure 5D-G). Here we have plotted the same data from Figure 4D into Figure 5D. Results for the first extinction session are described above, and detailed time-points for specific significance time windows can be found on Supplementary Table 1. On the last extinction session, we found increased LH-GABA activity after CS onset (Figure 5E).

Comparison between CS types in the last extinction session show that activity to the CS+ remained higher than CS- (Figure 5E). Comparison between the first and last extinction sessions revealed significant differences for CS+ (Figure 5F), but not CS- (Figure 5G). Overall, these data show that extinction is associated with both a decrease in alcohol seeking, and decreased cue-evoked activity of LH-GABA neurons.

#### Alcohol-priming induced reinstatement of extinguished alcohol seeking

We separated the animals into two groups: one group received primed 24 hour before the reinstatement with no priming on the reinstatement test (S5), and the other group received alcohol primed both 24 hours before and just prior to the reinstatement test (Figure 6)^31,32^ Only the second group showed reliable reinstatement to the CS+, and therefore we present these data here. Results for the first group are reported in the supplementary results (S4).

**Figure 6.**
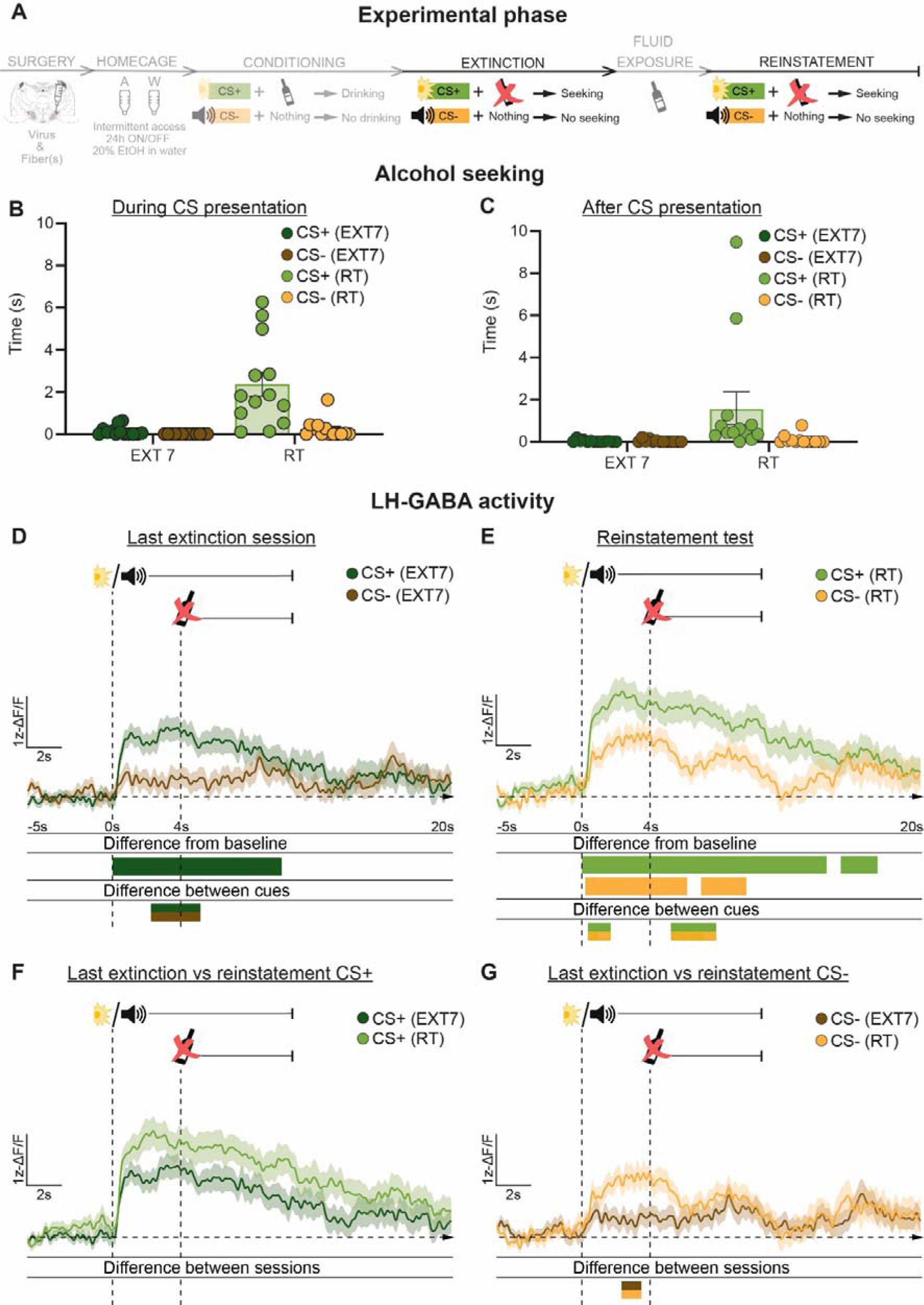
Monitoring LH-GABA during alcohol-priming induced reinstatement of alcohol seeking. (**A**) Outline of experimental procedure (n=13 females). Mean ± standard error of the mean (SEM) time spent in the alcohol magazine during (**B**) and after (**C**) CS presentation comparing the last extinction and reinstatement test sessions (n=8 trials per CS type, per session, individual data points show males (squares) and females (dots)). Ca2+ traces of LH-GABA activity centred around cue onset (-5s to +20s) comparing LH-GABA activity to CS+ and CS- on the last extinction (**D**) and reinstatement test (**E**) sessions. (CS+ EXT7: n=105; CS- EXT7: n=80; CS+ RT: n=92; CS- RT: n=73). The same mean Ca2+ traces are plotted for comparisons across sessions between CS+ (**F**) and CS- (**G**). For all photometry traces, bars at bottom of graph indicate significant deviations from baseline (dF/F ≠ 0), or significant differences between the specific events (CS+ EXT7; CS- EXT7; CS+ RT; CS- RT), determined via bootstrapped confidence intervals (99% CIs), and permutation tests with alpha 0.01 for comparisons between CS type and session. Vertical dashed lines indicate CS onset and alcohol delivery for CS+, horizontal line indicates baseline (dF/F=0).

<u>Behavioural data:</u> Repeated measures ANOVA comparing the last extinction session and reinstatement (Figure 6E and 6F) revealed a significant Session x CS Type interaction on time spent in the magazine during (Figure 6E; F(1, 12)=17,86, p < 0.01) but not after (Figure 6F; F(1, 12)=4,057; p=0.07) CS presentation. Specifically, we found that rats spent significantly more time in the alcohol port during CS+ on the reinstatement test, compared to during CS- presentation and during CS+ on the last extinction session (multiple comparisons, bonferroni corrected). These data show that the alcohol-prime procedure induced reinstatement of extinguished alcohol seeking, which was specific to the previously conditioned CS+. Repeated measures ANOVA comparing alcohol seeking across trials on the First Reinstatement session (n=8 trials per CS; Fig. S7B), revealed a significant Trial x Type of CS interaction both during (F(7, 84)=7,181, p < 0.001), and after (F(7, 84)=2,968, p < 0.01). This suggests behavioural reinstatement is extinguished within the first reinstatement session.

<u>Photometry data:</u> We made comparisons within a single session between the different CS types (CS+ vs CS-), and across the different sessions within the same CS type (Last Extinction vs. Reinstatement; Figure 6. D-G). Here we plotted extinction session data only for the reinstated group. Detailed time-points for specific significance time windows can be found on Supplementary Table 1. On the last extinction session (Figure 6D) we observed a significant increase from baseline for CS+ but not CS-. During reinstatement (Figure 6E), we found increased LH-GABA activity after CS onset for both cues. We found a significant difference in activity between CS+ and CS- on both the last extinction and reinstatement sessions. Despite the significant increase in alcohol seeking to the CS+ in reinstatement (Figure 6B), we found no significant differences in LH-GABA activity to CS+ between the last extinction session and reinstatement (Figure 6F), however we did observe a significant difference in CS- (Figure 6G).

We made comparisons between the first 4 and last 4 trials of reinstatement (Fig. S7D and Fig. S7F) and found significant differences in LH-GABA activity to the CS+, which was higher on the first 4 reinstatement trials compared to the last 4. This is also comparable to alcohol seeking, which we found to be higher in the first 4 trials compared to the last 4 (Fig. S7B).

### Exp 3. Optogenetic inhibition of LH-GABA impairs alcohol memory learning

Figures 7A and 7B show the experimental timeline. We used the dual-virus approach to confine expression of the inhibitory opsin GtACR2 (Mahn et al., 2018; Figure 1D-E) or control GFP/mCherry to LH-GABA neurons. We then trained the animals on the conditioning phase while delivering light to LH during presentation of both cues. We then tested the expression of the alcohol memory on a single extinction session, where no light inhibition or alcohol was delivered.

**Figure 7.**
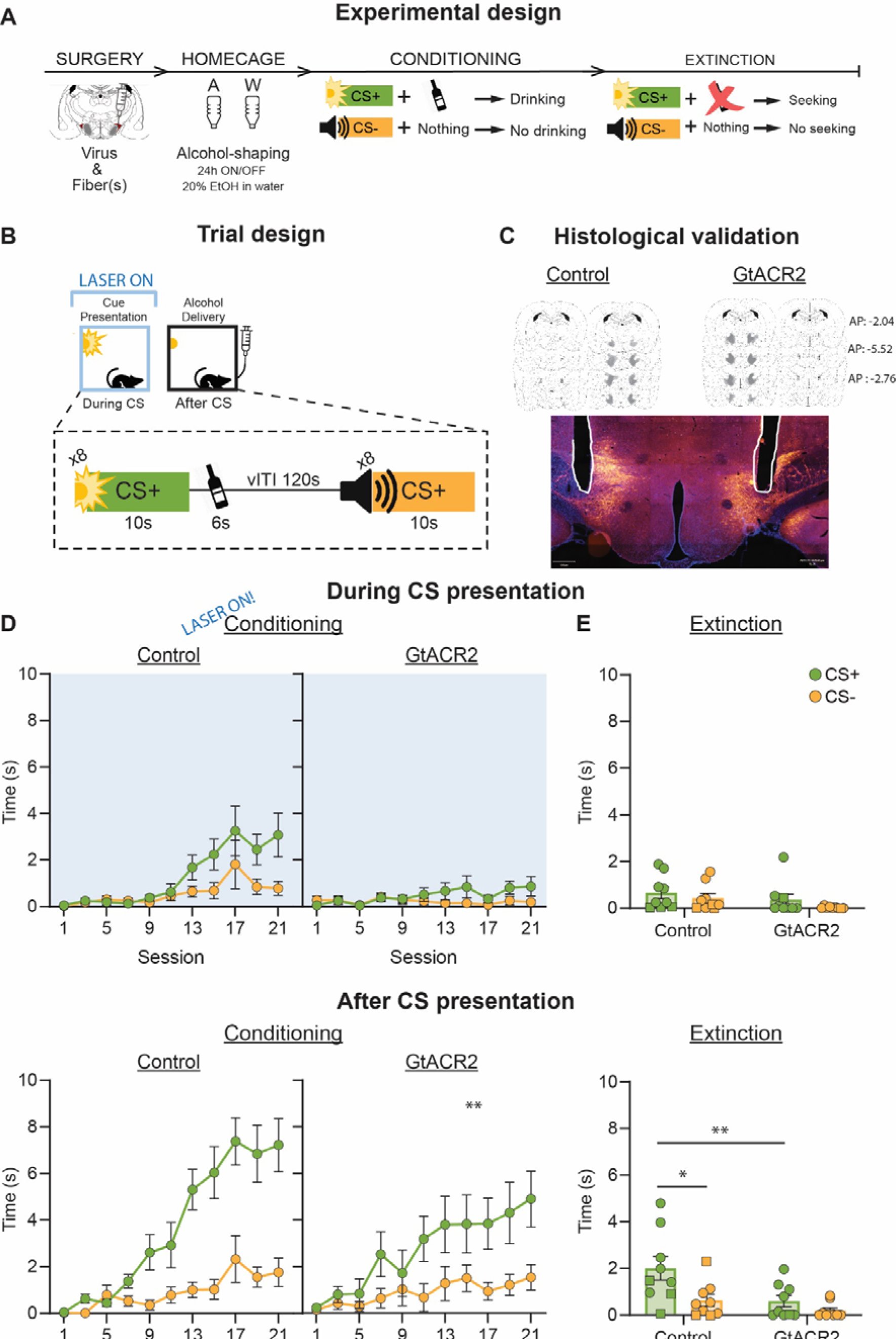
Optogenetic inhibition of LH-GABA during cue presentation in alcohol Pavlovian conditioning. (**A**) Outline of the experimental procedure (n=4 males, n= 14 females). (**B**) Trial design for optogenetics experiments. (**C**) Representative image of GtACR2 expression and fiber implants in LH, and fiber placement and expression validation. (**D**) Mean ± standard error of the mean (SEM) time spent in the alcohol magazine during (top) and after (bottom) CS presentation across conditioning sessions. Blue overlay depicts when optogenetic inhibition was delivered. (**E**) Mean ± standard error of the mean (SEM) time spent in the alcohol magazine during (top) and after (bottom) CS presentation on the extinction test. *p < 0.05.

Figure 7D shows alcohol seeking during (top; with light inactivation) and after (bottom) CS presentation throughout conditioning sessions. During CS presentation, we observed significant Session x CS Type (F(3,297, 52,75)=6,234, p < 0.001) and Session x Group (F(10, 160)=3,569, p >0.001) interactions, but no Session x Group x CS Type interaction.

Further analyses showed that the Session x Group interaction was only present in alcohol seeking to the CS+ (F(10, 160)=2,078, p < 0.05), but not CS-.

During the 10 seconds after CS presentation, we observed significant Session x Group (F(10, 160)=2,053, p < 0.95), Session x Type of CS (F(10, 160)=19,01, p < 0.001), and Session x Group x Type of CS (F(10, 160)=3,151, p < 0.01) interactions, but no Group x CS Type interaction. Further analyses showed that the Session x Group interaction was only present in alcohol seeking to the CS+ (F(10, 160)=2,944, p < 0.01), but not CS-. Overall, these data show that optogenetic inhibition of LH-GABA during conditioning prevented acquisition of the CS-alcohol memory.

Figure 7E shows alcohol seeking in the extinction test during (top) and after (bottom) CS presentation. We observed no significant effects of Group or CS Type, or an interaction between them during CS presentation. After CS presentation, we did observe a significant effect of Group (F(1, 16)=7,637, p < 0.05), as well as CS Type (F(1, 16)=8,959, p < 0.01), but no Group x CS Type interaction. Post-hoc analysis revealed a significant difference in alcohol seeking between groups to the CS+ (p < 0.001), but not the CS-. These data show that the control animals showed expression of the learned cue-alcohol memory, while the opsin group did not.

Finally, to rule out potential non-specific effects of our optogenetic manipulation on motivation to drink, rather than learning perse, we measured the amount of alcohol remaining at the end of each session and calculated total consumption in g/Kg (Fig. S2E). These data show that both the opsin and control group consume all alcohol in the sessions. Thus, optogenetic inhibition of LH-GABA during the cue period reduces learning about the cue but does not decrease alcohol consumption throughout the session.

## Discussion

In this study we describe the role of LH-GABA neurons in the acquisition and expression of a cue-alcohol memory. We first validated the dual-virus approach to confine Cre-dependent expression of fluorescent proteins, sensors, and opsins to LH-GABA neurons.

We then used this approach with fibre photometry calcium imaging in male and female rats to show that LH-GABA neurons increase activity to cues prior to learning, and as learning progressed the cue-evoked activity adapts. We found that LH-GABA activity is maintained for the alcohol-associated cue (CS+) but is decreased to the cue with no consequence (CS-). We also found evidence for prediction-error signalling during extinction, where LH-GABA neurons showed sustained activity when alcohol was expected but not delivered. Finally, during reinstatement we found that LH-GABA neurons maintain higher activity to the CS+ compared to the CS-, but this activity was not significantly higher than the last extinction session. Next, we used optogenetics in female rats to show that inhibition of LH-GABA during conditioning impaired learning of cue-alcohol associations. This study demonstrates the importance of LH-GABA in memory processes related to alcohol seeking.

### Methodological considerations

Several issues must be considered in the interpretation of these findings. The photometry and optogenetics experiments used slightly different conditioning protocol, where the alcohol was delivered during the cue in photometry but after the cue in optogenetics. We did this to avoid potential non-specific effects of LH-GABA inhibition on alcohol drinking. We found that alcohol delivery during the cue led to higher alcohol seeking during the cue period in experiment 2 compared to experiment 3. However, we also found that alcohol seeking during the post-cue period was not different between these experiments, providing evidence for learning in both training conditions.

This study describes learning associated with alcohol reward, but this does not necessarily relate to pathological alcohol intake as seen in alcohol use disorder. Because of the Pavlovian nature of our task, there is a limit to the amount of alcohol that can be consumed. However, the level of consumption we observed in the final sessions is approximately 0.9 g/Kg in a 45-minute session (Fig. S2D). While we did not measure blood alcohol, other studies have shown that this level of consumption in Long-Evans rats causes blood alcohol levels close to the binge level of 0.08 mg/dl^33^. It will be of interest in future studies to identify how LH-GABA activity relates to pathological alcohol intake.

In the reinstatement tests, we used two different procedures in two different cohorts of rats. In the first, the alcohol prime was given 24 hours prior to the test without alcohol primed during the reinstatement test^32^. In the second the alcohol prime was given both 24 hours before and at the beginning of the reinstatement test^34^. We found that the first group did not reliably reinstate alcohol seeking to the CS+, while the second group did. We therefore only analysed the reinstatement-photometry data from the second group.

The effect of sex and oestrous cycle has gained interest in recent years^35^. In all experiments, we used both male and female rats and did not observe any significant differences in behaviour or LH-GABA activity between the sexes. Therefore, we argue that the effect of opto-inhibition of LH-GABA neurons on acquisition of a cue-alcohol memory likely extends to both male and female rats.

Finally, although our findings show a specific role in cue-alcohol reward learning of the subpopulation of LH neurons defined by GAD expression, these are a heterogeneous population of neurons. This viral targeting strategy we used has been used in previous studies to restrict viral-mediated gene expression in GABAergic neurons in ventral tegmental area^29,36^. In LH, there is evidence from single-cell transcriptomics^37^ that GAD mRNA is expressed in all vGat (Slc32a1 mRNA) neurons, however this study also shows co-expression in glutamatergic neurons (Slc17a6 mRNA) in 4 of the 15 clusters they identified. Therefore, in this study we have recorded and manipulated activity of the LH-GABA neuron sub-population which is comprised of various phenotypes and efferent targets. It will be of interest in future studies to determine whether sub-populations of GAD+ neurons differentially contribute to the effects we have observed here.

Cue-induced activity of LH-GABA neurons changes with acquisition of a cue-alcohol association and modulate their activity across extinction learning.

Throughout all phases of the experiment, we found LH-GABA activity is increased to the presentation of a cue. This observation shows that LH-GABA activity changes in response to most changes in the environment. Interestingly there is precedent for a function such as this from historical lesion data, showing sensory neglect from LH lesions^38,39,40^. However, an important finding of this study learning is associated with significant changes in the patterns of activity.

On the first conditioning session, we found LH-GABA activity increases to both cues which initially have no motivational relevance. By the last conditioning day, LH-GABA activity was significantly higher to CS+ compared to CS-. Thus, we show that LH-GABA activity significantly decreased to the non-predictive cue and is sustained to the alcohol-predictive cue. This pattern of activity might reflect a role of LH-GABA in gating activity to motivationally significant stimuli, maintaining activity to behaviourally relevant stimuli, while decreasing activity to irrelevant ones. This function may be consistent with observations that LH is important for attentional processes involved in learning^41^. Consistent with this, on the first extinction session we found that activity to the alcohol-predictive cue remained significantly higher compared to the last conditioning session after the cue was turned off. This sustained activity might indicate that LH-GABA is signalling an expectation for alcohol delivery. This signal could contribute to a reward prediction error signal. Reward prediction errors signal violation of an expected reward^42,43^, and we have previously proposed that LH-GABA projections to VTA are involved in the generation of the prediction error signal in VTA dopamine neurons^17^. Furthermore, violations in expected outcomes also increase attentional processes^44^, providing further evidence for a role in LH-GABA in attentional processes.

In the last extinction session, we found that LH-GABA activity to the CS+ is significantly decreased compared to the first extinction session. We propose that this further demonstrates a role of LH-GABA in acting as an information gating filter^19^. In this model, LH-GABA activity is decreased to the CS+ because extinction has weakened the incentive value of this cue. Interestingly, even after alcohol seeking is fully extinguished, we still observe significant differences in LH-GABA activity between the CS+ and CS-. This suggests that LH-GABA activity to the formerly alcohol-predictive cue is not completely erased by extinction, which is consistent with the view that extinction is not memory erasure but rather creates a new memory which gates the expression of the originally learned memory^45,46^.

Finally, during reinstatement we found some evidence that LH-GABA activity to the alcohol-predictive CS+ is increased, specifically in the earlier trials, indicating that the incentive value of the cue can be retained after extinction.

These processes may be orchestrated through excitatory projections from the basolateral amygdala (BLA) to the LH^47^, which are known to signal salient sensory-specific properties of stimuli based^44,48,49^. Given the strong reciprocal projections of LH-GABA and VTA^50,51,24^, and the role this area is suggested to play in attention^41^, a key component of associative learning^52^, and memory updating^19^, it is likely that LH-GABA play a role in increasing attention to encode new stimulus-outcome associations.

Potential contribution of LH-GABA to alcohol priming-induced reinstatement.

During priming-induced reinstatement, we found that LH-GABA activity to the CS+ was significantly higher than to the CS- for some periods of the cue presentation. However, activity to the CS+ was not increased relative to the last extinction session. When comparing the reinstated (Fig. 6) with the non-reinstated (Fig. S5), only the reinstated group shows significantly higher activity to the CS+ compared to the CS-, which was not present in the non-reinstated group. Furthermore, within-session analysis (Fig. S7) of the LH-GABA activity during reinstatement shows that cue-induced activity to the CS+ was higher in the first 4 trials than the last 4. As such, our data do point to a potential role for LH-GABA neuronal activity in alcohol priming-induced reinstatement of extinguished alcohol seeking.

This suggests that the function of LH-GABA might extend to reinstatement by increasing their activity to the cues when these again become motivationally relevant. The LH has previously been implicated in reinstatement procedures^53^, and addictive behaviours are characterised by attentional biases for drug-related cues^54,55^. Given the LH role in attentional processes this area could be involved during relapse by directing attention to drug related cues.

### Optogenetic inhibition of LH-GABA impairs learning of alcohol-predictive cues

In experiment 3 we show that optogenetic inhibition of LH-GABA neurons during cue presentation significantly impaired learning of the alcohol-predictive cue. This observation is consistent with the photometry observations during conditioning. These conclusions were strengthened during the extinction test without opto-inhibition of LH-GABA neurons, where only the control group showed alcohol seeking post CS+. This observation is comparable to our previous work in the transgenic GAD-Cre rat, showing that opto-inhibition of LH-GABA neurons impairs acquisition of a cue-food association^17^, and thus extends the role of LH-GABA neurons in reward learning to addictive substances such as alcohol.

The LH harbours a wide range of different neuropeptide expressing neurons, such as Orexin/hypocretin (Ox/Hcrt^56^), Melanin-Concentrating Hormone^56^ (MCH) Neurotensin, Galanin^41,16^, as well as putative γ-aminobutyric acid- (GABA) and Glutamate- releasing neurons^58^. These diverse neuronal sub-populations have been shown to host a variety of functions, including associated with alcohol seeking^59,60^ and associative learning^61,62^. The contribution of LH-GABA to these functions may be consistent with the findings from these studies, indicating that the multiple different cellular phenotypes within LH contribute towards addictive behaviours^53^.

## Concluding remarks

In this study we sought to investigate the role of LH-GABA in learning about stimuli which predict alcohol rewards. Our results show that activity in LH-GABA is necessary for the acquisition of alcohol-predictive cues. We also show that the activity of LH-GABA neurons is modulated through experience, based on changes in the motivational properties of stimuli across learning. Future studies are needed to determine the extent to which this activity is necessary for relapse, by manipulating these neurons activity during reinstatement procedures. Overall, our findings highlight an important role for LH-GABA in a fundamental learning process and show that this system is a critical component of memory formation in alcohol addiction.

## Supporting information

Supplementary Results

## Acknowledgements

The work was supported by an NWO VIDI grant (016.Vidi.188.022).

The authors gratefully acknowledge the VUmc Histology Imaging Unit for their support & assistance in whole-slide imaging. The authors gratefully acknowledge Dr. Philip Jean-Richard-Dit-Bressel for assistance with photometry analysis and sharing early versions of the scripts we used.

## Author contributions

Conducted experiments: IAL, DS, YvM, ST, PW, LA, CvdN, TH. Analysed data: IAL, DS, ST, PW, LA, CvdN, TH. First draft of manuscript: IAL, NJM. Edited subsequent drafts and finalised manuscript: IAL, NJM, AJM, DS, YvM; TdV, HM.

## STAR Methods

### RESOURCE AVAILABILITY

#### Lead contact

The lead contact for this study is Dr. Nathan J. Marchant.

#### Materials availability

This study did not generate new unique reagents. Further information and requests for resources and reagents should be directed to and will be fulfilled by the Lead Contact.

#### Data and code availability

- All data reported in this paper will be shared by the lead contact upon request.
- All original code has been published at <u>github.com/ialozares/AlcoholConditioning</u> and is publicly available as of the date of publication.
- Any additional information required to reanalyze the data reported in this paper is available from the lead contact upon request.

### EXPERIMENTAL MODEL AND STUDY PARTICIPANT DETAILS

We used 40 adult wild-type Long-Evans rats (8 males and 32 females), from Janvier (France). All behavioural and surgical procedures were performed in the animal facility of the Vrije Universiteit, Amsterdam, The Netherlands. Rats were kept in temperature and humidity-controlled rooms, in an inverted 12/12 light/dark cycle, and we trained the rats during the dark cycle. Food and water were provided ad libitum. Animals were kept in cages of either 2 or 3 cage mates. All procedures were approved by the Instantie voor Dierenwelzijn of the Vrije Universiteit, Amsterdam, The Netherlands.

We did not make specific analysis to determine sample size prior to any experiments.

Cohorts were balanced by experimental group prior to the experiment starting. We allocated the rats randomly to one of the groups within the optogenetics and chemogenetics experiments. We did not exclude any rats for reasons of behavioural validation. Rats in Experiments 2 and 3 that had expression or fiber placement not within LH, were excluded form analysis. Rats in Experiment 3 that had either unilateral expression, misplaced expression or a misplaced fiber placement were excluded from analysis.

### METHOD DETAILS

#### Apparatus

All procedures were performed in standard Med Associates operant chambers with data collected through the MED-PC V program (Med Associates, Georgia, VT, USA). Each chamber had a grid floor and a large opening at the top to allow patch cords through. Each chamber was equipped with a white noise generator, and one or two light cues, as well as a magazine port where the alcohol was delivered (20% ethanol). The magazines were equipped with a beambreak device to measure total time spent in the magazine. Custom made Med-PC programs controlled all aspects of the task.

For fiber photometry, excitation and emission light was relayed to and from the animal via optical fiber patch cords (0.48 NA, 400-µm flat tip; Doric Lenses). Blue excitation light (490 or 470 nm LED [M490F2 or M470F2, Thorlabs]) was modulated at 211 Hz and passed through a 460–490 nm filter (Doric Lenses), while isosbestic light (405 nm LED [M405F1, Thorlabs]) was modulated at 531 Hz and passed through a filter cube (Doric Lenses).

GCaMP7f fluorescence was passed through a 500- to 550-nm emission filter (Doric Lenses) onto the photoreceiver (Newport 2151). Light intensity at the tip of the fiber was measured before every training session and kept at 21 µW. A real-time processor (RZ5P, Tucker Davis Technologies) controlled excitation lights, demodulated fluorescence signals and received timestamps of behavioural events. Data were saved at 1017.25 Hz and analysed with custom-made Matlab scripts, available at: https://github.com/ialozares/AlcoholConditioning

For optogenetics experiments, light (470nm) was delivered to the animals via optical bifurcating fiber patch cords (0.48 NA, 200-µm flat tip; Doric Lenses) connected to the head of the animals. Light delivery began 500ms prior to CS onset until 500ms after CS presentation. Light intensity was kept at 5mW and measured at the beginning of each session for each operant chamber. Light delivery was controlled by custom Med-PC scripts. **Drugs**

Deschloroclozapine (DCZ) (MedChem Express, 1977-07-7) was dissolved in 1-2% dimethyl sulfoxide (DMSO) in saline to a final volume of 0.1ml per kg. Alcohol solutions were prepared using 96% (v/v) ethanol diluted with tap water and were administered using normal bottles.

#### Surgeries

The day before and thirty minutes prior to surgery, we injected rats with the analgesic Rymadil® (5 mg/kg; Merial, Velserbroek, The Netherlands) Surgery was performed under isoflurane gas anesthesia (PCH; Haarlem). The surgical area on the skull was shaved & disinfected before surgery and the skin was locally anesthetized using a subcutaneous injection of 0.1 ml 10% lidocaine. Craniotomies above LH, VTA or NaCs were performed, followed by AAV injections using a micro-syringe connected to a micropump (UltraMicroPump 4, WPI surgical instruments), which was lowered into the brain until it reached the DV coordinates (see below for details). Additionally, for fibre photometry and optogenetics experiments, 6 screws were drilled on the skull to support the cement needed for fibre anchoring. Next, the optic fibres were implanted using the same stereotactic apparatus and bregma-relative coordinates as used for the AAV injection. The optic fiber implant(s) (when applicable) were then secured to the skull using dental cement (IV Tetric EvoFlow 2g A1, Henry Schein, Almere) and jewellers screws. Rymadil (5 mg/kg; s.c.) was administered for 2 days after the surgery. Rats were given one week of recovery following surgery.

#### AAV combinations

For all experiments, we injected a cocktail of one AAV expressing Cre under the GAD promotor, and another AAV encoding Cre-dependent jGCamP7f, GTACR2, or control GFP or mCherry. For unilateral inhibition experiments, we additionally injected an AAV-retro encoding flp in LH, and an AAV encoding FLP-hm4di or control mCherry.

#### AAV injections for validation experiments

1µl of AAV solution was injected bilaterally into LH (AP: + 2.3, ML: 2, DV: -8.4 to -8, two infusions) over 2.5 min, the needle was then left in place for 5 min, brought up to DV:-8, and then injected again for 2.5 min and left to diffuse for 5 min.

#### AAV injections for fiber photometry

1µl of AAV solution was injected unilaterally into LH (AP: + 2.3, ML: 2, DV: -8.4 to -8, two infusions) over 2.5 min, the needle was then left in place for 5 min, brought up to DV:-8, and then injected again for 2.5 min and left in place for 5 min. A 400µ optic fiber (Doric Lenses) was then implanted above LH.

#### AAV injections for optogenetics

0.5µl of AAV solution was injected unilaterally into LH (AP: + 2.3, ML: 2, DV: -8.4 to -8, two infusions) over 2.5 min, the needle was then left in place for 5 min, brought up to DV:-8, and then injected again for 2.5 min and left in place for 5 min. A 200µ optic fiber (Doric Lenses) was then implanted above LH.

#### AAV injections for chemogenetics

0.5µl of AAV solution was injected unilaterally into either NAcS(AP: + 1.7, ML: + 2.4, DV: -7.6) or VTA (AP: -5.4, ML: + 2.61, DV: -7.5) over 5min. The needle was then left in place for 5 min.

#### Behavioural procedures

##### Intermittent access homecage alcohol

For all behavioural experiments, we first habituated rats to alcohol consumption by using an alcohol homecage procedure (Simms et al., 2008; Wise RA, 1973; Supp. Fig. 1). Rats had intermittent 24h access to a bottle containing 20% ethanol (EtOH), and a bottle containing water. Additionally, we used two control empty cages without animals to account for spillage. After 24h the bottles were weighed and swapped for a single water bottle for the following 24h. These two phases were repeated 12 times. The overall consumption of alcohol per rat after each session was measured in grams, by subtracting pre and post measurements for each cage and the average of the two control cages, accounting for 2- gram spillage and multiplying by the density of 20% ethanol (0.97).

During this phase, we habituated the rats to the optogenetics or photometry patch cords by tethering them to the patchcord and leaving them in their designated behavioural boxes for 30 minutes on the alcohol off days (Tuesday and Thursday). Alcohol magazines were removed from the boxes during habituation. For the chemogenetics inhibition experiment, we habituated the rats to an i.p. Injection by injecting approx. 0.1ml saline i.p. prior to 4 habituation sessions.

##### Alcohol Pavlovian conditioning task

**Conditioning Phase:** We placed the rats in the operant chambers described earlier and trained once a day during weekdays (Mon-Fri) for 45 minutes per group, per day. In these sessions, we trained the rats to associate a cue with alcohol delivery in the alcohol magazine (CS+ trials) and a different cue with no programmed consequences (CS- trials). Cues consisted of three different stimuli (white noise, light, or flickering light) counterbalanced across rats. In each session, rats received 8 presentations of each CS (16 trials total). Trials were further divided into three phases (pre-CS, during-CS, and post-CS), each lasting 10 seconds and resulting in a 30-second trial window. Alcohol (20% EtOH) was delivered for 6 seconds into the magazine for CS+ trials (0.2ml per trial, for a total of 1.6ml each session).

Trials were followed by a variable inter-trial-interval (vITI 2mins). Drinking was assessed by measuring the time beam breaks in the magazines were broken for all trial and vITI phases. Alcohol leftover was measured after each training session to assess the amount animals were drinking. For all experiments, we gave every rat between 19-21 conditioning sessions.

#### Extinction phase

We placed the rats in the operant chambers and trained them once a day during weekdays for 45 minutes per day. Extinction sessions were procedurally identical to the conditioning sessions, with the exception that no alcohol was delivered after the CS+. Alcohol seeking was again assessed by measuring the time spent in the magazine during all CS trial and vITI phases.

#### Fluid exposure session

Rats received a single fluid exposure session after the last extinction session. In this session, alcohol was delivered in the alcohol magazine on the same time schedule as the conditioning phase, but the cues were not presented.

#### Alcohol-prime induced reinstatement test

Alcohol was delivered into the magazine for 6 seconds at the beginning of the test session. During the test session, both cues were presented in a similar manner to the conditioning and extinction sessions, but no alcohol was delivered. Alcohol seeking was measured as the time spent in the magazine during all CS and vITI phases.

#### Histological validation

##### Immunohistochemical labelling of MCH and Ox

After a waiting period of approximately 4 weeks to allow for virus expression, we deeply anesthetized rats with isoflurane and Euthasol® injection (i.p.) and transcardially perfused them with ∼100 ml of normal saline followed by ∼400 ml of 4% paraformaldehyde in 0.1M sodium phosphate (pH 7.4). The brains were removed and post-fixed for 2 h, and then 30% sucrose in 0.1M PBS for 48 h at 4°C. Brains were then frozen on dry ice, and coronal sections were cut (40 µm) using a Leica Microsystems cryostat and stored frozen in 0.1M PBS containing 30% sucrose at -20°C.

We selected a 1-in-4 series and first rinsed free-floating sections (3 x 10 minutes) before incubation in PBS containing 0.5% Triton-X and 10% Normal Donkey Serum (NDS) and incubated for at least 48 h at 4°C in primary antibody. Sections were then repeatedly washed with PBS and incubated for 2-4 h in PBS + 0.5% Triton-X with 2% NDS and secondary antibody. After another series of washes in PBS, slices were stained with DAPI (0.1 ug/ml) for 10 min prior to mounting onto gelatin-coated glass slides, air-drying and cover-slipping with Mowiol and DABCO.

Primary antibodies were guinea pig anti-orexin (1:250; Synapic Systems ART: 389004), and rabbit anti-MCH (1:500; Phoenix Biotech ART: G-070-47). Secondary antibodies were Alexa Fluor 488 anti-guinea pig (1:500; Molecular Probes ART: A11073) and IRDye 680LT anti-rabbit (1:500; Li-cor ART: 926-68023).

#### In-situ hybridization labelling of GAD-mRNA

We performed RNA in situ hybridization for glutamate decarboxylase1 (GAD1) and eGFP mRNA and according to “User Manual for Fresh Frozen Tissue’’ from RNAscope Multiplex Fluorescent Reagent Kit V2 assay (Advanced Cell Diagnostics). Specifically, freshly frozen brains were cryosectioned (20 μm) onto Superfrost Plus glass slides (Fisher Scientific) and stored at −80°C. Brain sections were fixed in 10% neutral buffered formalin for 15 min at 4°C, rinsed twice in distilled water, gradually dehydrated for 5 min each in 50%, 70% and twice 100% ethanol. The slides were then incubated with Hydrogen Peroxide for 10 min, and Protease IV for 30 min at RT. In between steps sections were rinsed twice in 1xPBS. After this, GAD1 (316401-C2) and eGFP (400281) specific target probes were applied to the sections and incubated at 40°C for 2 hr in the HybEZ oven (Advanced Cell Diagnostics).

Sections were then treated with amplifier probes by applying AMP1 and AMP2 at 40°C for 30 min each, followed by AMP3 at 40°C for 15minfor 30 min. Signals for channel 1 (eGFP, Opal520), and channel 2 (GAD, Opal690) were developed by applying the appropriate HRP probes onto the tissue. Finally, the nuclei were stained using DAPI for 30 s to stain nuclei (blue color). In between steps sections were rinsed twice in 1x Wash Buffer (Advanced Cell Diagnostics).

#### Histological validation after behavioural experiments

After each behavioural experiment, we deeply anesthetized rats with isoflurane and Euthasol® injection (i.p.) and transcardially perfused them with ∼100 ml of normal saline followed by ∼400 ml of 4% paraformaldehyde in 0.1M sodium phosphate (pH 7.4). The brains were removed and post-fixed for 2 h, and then 30% sucrose in 0.1M PBS for 48 h at 4°C. Brains were then frozen on dry ice, and coronal sections were cut (40 µm) using a Leica Microsystems cryostat and stored frozen in 0.1M PBS containing 30% sucrose at - 20°C.

We selected a 1-in-4 series and first rinsed free-floating sections (3 x 10 minutes) before washing in PBS. Following this, slices were stained with DAPI (0.1 ug/ml) for 10 min prior to mounting onto gelatin-coated glass slides, air-drying and cover-slipping with Mowiol and DABCO.

#### Image acquisition and neuronal quantification

We used a VectraPolaris slide scanner (VUmc imaging core) to image slides at 10x magnification. For validation of GAD-cre approach experiments (Exp. 1), images from Bregma -2.04 to Bregma -2.76 were scanned and imported into QuPath for analysis. For photometry and optogenetic experiments, images containing LH, from Bregma - 2.04 mm to -2.76 mm were identified and the boundary of expression and fiber placement for each rat was plotted onto the respective Paxinos and Watson atlas. For chemogenetic experiments images from Bregma 2.16 to Bregma 1.56 (NAcS), and Bregma -4.92 to Bregma -5.40 (VTA) were identified and the boundary of expression was plotted onto the respective Paxinos and Watson atlas.

LH was manually labelled using DAPI for identification of anatomical landmarks and boundaries. Each value was the result of an average of each count from 4-5 adjacent sections per rat (approx Bregma -2.04 mm to 2.76 mm). To identify GAD-, eGFP-, MCH-, Ox-, and redFusion-positive cells, we used the ‘Cell detection’ feature in QuPath, with an identical threshold applied across all sections. To identify GAD+eGFP, MCH+eGFP(or redFusion), and OX+eGFP(or redFusion), each region of interest was exported to ImageJ. The overlays representing the cells (GAD, eGFP, MCH, Ox, and redFusion) were then filled, converted to a binary layer, and then multiplied using the ImageJ function ‘Image calculator’. Double labelling is reported as a percentage of total neurons labelled with the AAV fluorescent reporter for that given region of interest.

#### Ex vivo slice physiology

Rats were anesthetized (5% isoflurane, i.p. injection of 0.1 ml/g pentobarbital) and perfused with ice-cold N-methyl-D-glucamin (NMDG) solution containing (in mM): 93 NMDG, 2.5 KCl, 1.2 NaH2PO4, 30 NaHCO3, 20 4-(2-hydroxyethyl)-1-piperazineethanesulfonic acid (HEPES), 25 glucose, 5 sodium ascorbate, 3 sodium pyruvate, 10 MgSO4, 0.5 CaCl2, at pH 7.3 adjusted with HCl. After removing the brain, 250 μm thick brain slices were cut in ice-cold NMDG solution using a Leica VT1200 vibratome. The slices were transferred to a holding chamber with carbogenated artificial cerebrospinal fluid (aCSF; in mM: 125 NaCl, 3 KCl, 1.2 NaH2PO4, 1 MgSO4, 2 CaCl2, 26 NaHCO3, and 10 glucose). Slices were stored at room temperature for at least 1 hour before transferring them to the recording chamber.

During recordings slices were kept at 34°C in aCSF. Whole cell recordings were made using borosilicate glass patch-pipettes (3–5 MΩ) were used with a K-gluconate-based internal solution containing (in mM): 135 K-gluconate, 4 NaCl, 2 MgATP, 10 phosphocreatine, 0.3 GTP (sodium salt), 0.2 EGTA, 10 HEPES at a pH of 7.3 using a Multiclamp 700B amplifier (Molecular Devices) and pClamp software (Molecular Devices). Optogenetic stimulation was done at 470 nm using a DC4100 4-channel LED-driver (Thorlabs, Newton, NJ) as light source.

#### Experimental design

Exp 2.a: Monitoring LH-GABA neurons during alcohol pavlovian conditioning (n=3 female, n=2 male)

Fig 2A shows the experimental outline. Alcohol delivery coincided with presentation of the CS+, beginning 4 seconds after CS+ onset. All phases were carried out as described in the behavioural procedure.

Exp 2.b: Unilateral chemogenetic inhibition and monitoring neurons during alcohol Pavlovian conditioning (n=13, female)

Supp. Fig. 4 shows the experimental outline. In this experiment we delivered 0.1 mL of 20% alcohol into the magazine. Alcohol delivery coincided with presentation of the CS+, beginning 4 seconds after CS+ onset. All phases were carried out as described in the behavioural procedure. We used chemogenetics to chemogenetically inactivated projections from VTA or NAcS to LH on the days 1 or 2, and 17 or 18 (counterbalanced d-clz or sal) of the conditioning phases, and on the first extinction day. For tests during other conditioning phase, we injected both GFP and hM4D9i expressing rats with d-clz (0.1 mg/kg injection i.p.) or saline (0.1 mg/kg injection i.p), counterbalanced across the two test days. For the first extinction day test, we injected all animals with d-clz (0.1 mg/kg) before the start of the session. We found very sparse or off target DREADDs expression (Supp. Fig. 3), leading to sample sizes not big enough to make statistical comparisons. Moreover, we found no d-CLZ effects on LH-GABA activity or behaviour (Supp. Fig. 4) and given the behavioural task was identical, we combined behavioural and neural data from animals in experiments 2a and 2b. <u>Exp 3: Inhibiting LH-GABA neurons during encoding of alcohol-predictive cues (n=4 males,</u> <u>n=14 females)</u>

Fig 7A shows the experimental outline. During the presentation of the CS, light (470 nm) was delivered into the LH, thereby inhibiting the GABAergic neurons. Light delivery occurred via a bifurcating patch-cord (Doric Lenses Inc) connected to the optic fibres on the head of the rat. Light delivery occurred only during the conditioning phase, and alcohol was delivered 0.5 seconds after CS+ offset. This was done to avoid potential effects of LH-GABA inhibition on drinking. After conditioning, we tested all rats in a single extinction session. During this session, cues were presented in the same manner as during conditioning, but there was no light delivery during the cue presentation, and no alcohol delivered after CS+ offset.

### QUANTIFICATION AND STATISTICAL ANALYSIS

Behavioural data were analysed using GraphPad Prism 9.1.0. Phases were analysed separately. Dependent variables were the averaged time spent in the magazine during CS+ and during CS-, or post CS+ and post CS- per session across phases (Conditioning, Extinction, Reinstatement).

#### Experiment 2 - Behaviour

For the first Extinction and first Reinstatement sessions, we used a repeated measures analysis of variance (ANOVA) with type of CS (CS+, CS-), and trial number (Trial 1 – Trial 8) as within-subjects factors to assess changes in alcohol seeking within session. For all other analyses, we used a repeated measures analysis of variance (ANOVA), with type of CS (CS+, CS-) and session (Across Conditioning sessions, Last Conditioning vs. First Extinction; Across Extinction sessions; Last Extinction vs Reinstatement) as the within-subject factors.

#### Experiment 2 - Photometry

Recorded signals were first down sampled by a factor of 64, giving a final sampling rate of 15.89 Hz. The 405 nm isosbestic signal was fit to the 490 nm calcium-dependent signal using a first-order polynomial regression. A normalized, motion-artifact-corrected ΔF/F was then calculated as follows: ΔF/F=(490 nm signal − fitted 405 nm signal)/fitted 405 nm signal. The resulting ΔF/F was then detrended via a 90-s moving average, and low-pass filtered at 3 Hz. ΔF/F from 10 s before CS (baseline) to 20 s after CS onset were collated. These traces were then baseline corrected and converted into z-scores by subtracting the mean baseline activity during the first 5 s of the baseline and dividing by the standard deviation of those 5 s.

Traces were grouped by type of CS (CS+ or CS-) and by session (First conditioning, Last Conditioning, First Extinction, Last Extinction, Reinstatement Test). For within session analyses on the First Extinction and Reinstatement sessions, traces were grouped by first 4 and last 4 trials, reflecting early and late extinction and reinstatement. Two analysis approaches were used, bootstrapping and permutation tests, the rationale for each is described in detail in Jean-Richard-Dit-Bressel et al., 2020 and Garreh et al., 2022. The output of every statistical test we conducted is available in the raw data files.

Bootstrapping was used to determine whether calcium activity type of CS was significantly different from baseline (ΔF/F=0). A distribution of bootstrapped means were obtained by randomly sampling from traces with replacement (n traces for that response type; 5000 iterations). A 95% confidence interval was obtained from the 2.5th and 97.5th percentiles of the bootstrap distribution, which was then expanded by a factor of sqrt (n/(n − 1)) to account for narrowness bias (Jean-Richard-Dit-Bressel et al., 2020).

Permutation tests were used to assess significant differences in calcium activity between CS+ and CS- across the different phases (First Conditioning, Last Conditioning, First Extinction, Last Extinction, and Reinstatement sessions) of the task. Observed differences between CS type or session were compared against a distribution of 1000 random permutations (difference between randomly regrouped traces) to obtain a p value per time point. Alpha of 0.05 was Bonferroni corrected based on the number of comparison conditions, resulting in alpha of 0.01. For both bootstrap and permutation tests, only periods that were continuously significant for at least 0.25 s were identified as significant (Jean-Richard-Dit-Bressel et al., 2020).

The specific time windows of significance from these two statistical tests are described in Supplementary Table 1.

#### Experiment 3 - Behaviour

We used a 3-way ANOVA to determine the effects of group (Control vs. Experimental), type of CS (CS+, CS-), and session (Across Conditioning sessions, Last Conditioning vs. First Extinction) and the interaction between them.

