## Supplementary Results for "Lateral hypothalamic GABAergic neurons encode alcohol memories"

**Exp. 2b: Unilateral chemogenetic inhibition and monitoring neurons during alcohol Pavlovian conditioning**

In experiment 2b., we aimed to unilaterally inhibit VTA and NAcS projections to LH-GABA using DREADDs. We used fiber photometry to measure real-time population level LH-GABA neuron calcium (Ca2+) transients on test days. Supplementary Figure 3 shows histological validation of the viral approach, and Supplementary Figure 4 shows a lack of effect of unilateral inhibition on both alcohol seeking and LH-GABA calcium activity. Validation of the expression of the viral constructs used to target VTA and NAcS projections to LH-GABA indicated insufficient expression, as such we combined both groups determine whether there were any non-specific effects of the chemogenetic ligand on alcohol seeking or LH-GABA calcium activity.

Behavioural data: We used repeated measures ANOVA with the within-subjects factors Drug (Saline, d-clz) and CS Type (CS+, CS-) to compare alcohol seeking during the vehicle and d-clz sessions on the First (Supp. Fig. 4, left), Last (Supp. Fig. 4C, left) Conditioning and Extinction (Supp. Fig. 4D, left) sessions. We found no effect of Drug on alcohol seeking. On the first conditioning tests we found no effect of Drug or CS Type. On the last conditioning tests we found a main effect of CS Type (During: (F (1, 12) = 60.41, p < 0.0001; After: F(1, 12) = 12.62, p < 0.001), however we found no effect of Drug, indicating that neither the ligand nor the chemogenetic inhibition caused a change in alcohol seeking. Finally, although the data reflects an effect of CS type (During: F (1, 12) = 13,00, p < 0.01; After: F (1, 12) = 25,62, p < 0.001) and Test (During: F (1, 12) = 12,99, p < 0.01; After: F (1, 12) = 11,84, p < 0.01), this effect is not related to the manipulation, and reflects normal extinction learning since we observe extinguished alcohol seeking already in the second session.

Photometry data: We made comparisons between the different CS Type (CS+ vs CS-), and across the different test sessions within the same CS type (Saline, D-CLZ). On the First Conditioning Tests (Supp. Fig. 4B, right), bootstrapping analysis revealed a significant increase from baseline for CS+ and CS- on the Vehicle and D-CLZ tests. Permutation analysis revealed no significant differences.

On the Last conditioning tests (Supp. Fig. 4C, right), bootstrapping analysis revealed a significant increase from baseline for CS+ and CS- on the Vehicle and D-CLZ tests. Permutation analysis revealed significant differences between CS+ and CS-. We found no significant differences between test sessions.

On the Extinctions tests (Supp. Fig. 4D, right), bootstrapping analysis revealed a significant increase from baseline for CS+ and CS- on the Vehicle and D-CLZ tests. Permutation analysis revealed significant differences between CS+ and CS-. We found no significant differences between test sessions.

These results show that the chemogenetic manipulation had no effect on alcohol seeking or on LH-GABA calcium activity. This is supported by the lack of expression of the viral constructs. As a result, we combined the behavioural and photometry data from experiments 2a and 2b.

**Supplementary Figures**

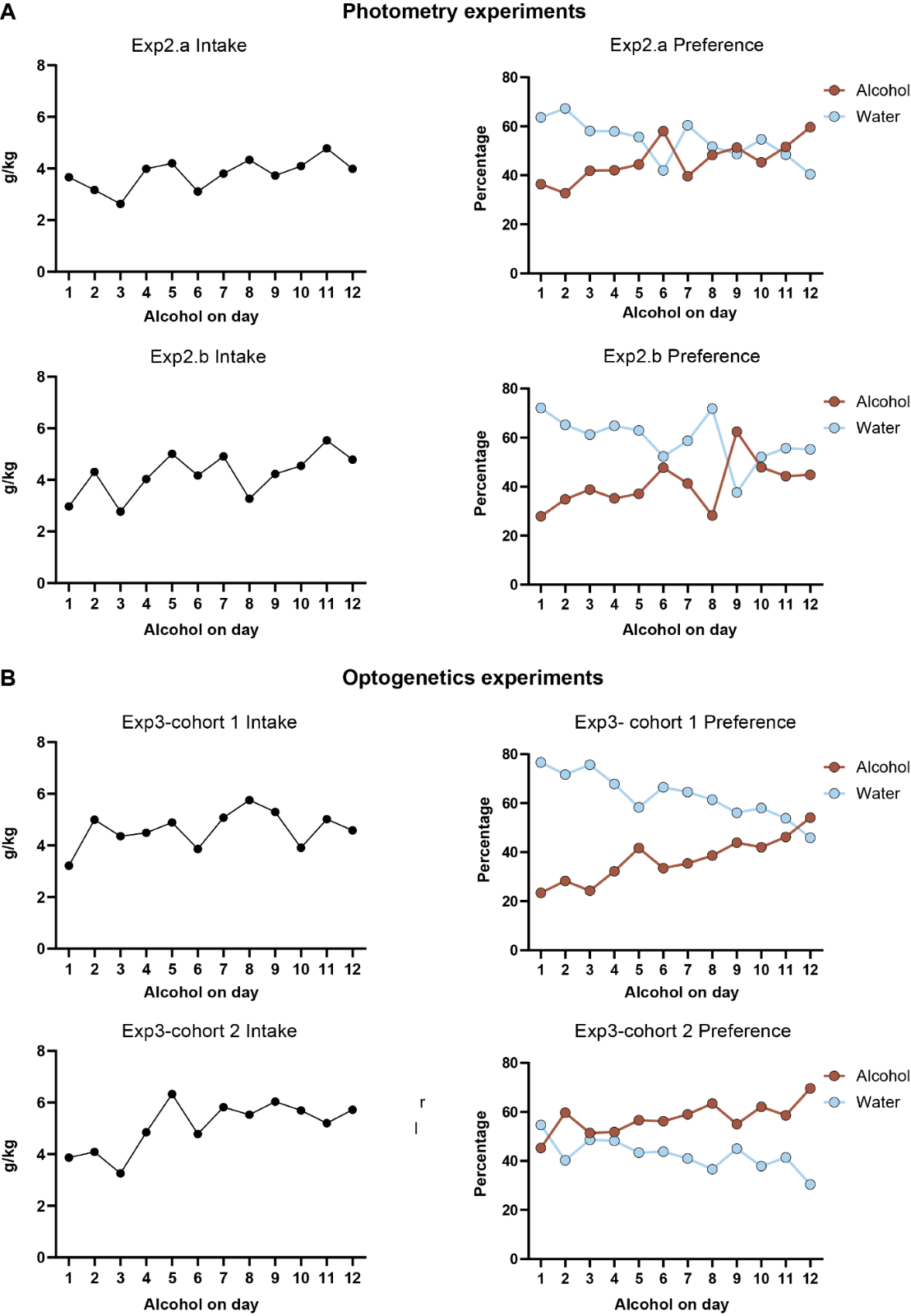

**Figure S1. Alcohol-home cage intake and preference**

Mean consumption (left) and preference (right) of alcohol and water across homecage sessions for photometry **(A)** and optogenetics **(B)** experiments.

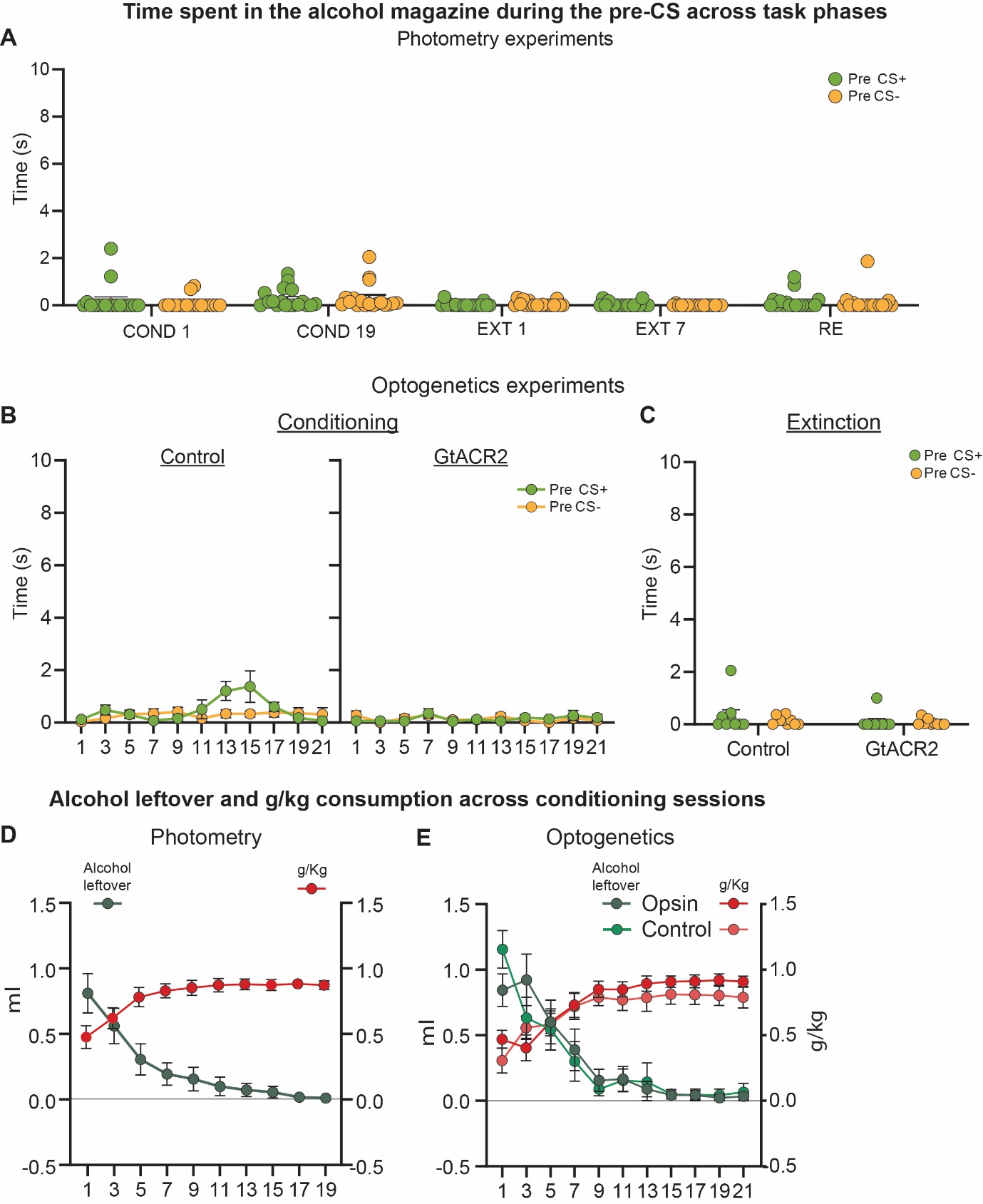

**Figure S2. Time spent in the alcohol magazine during pre-CS period, and alcohol consumption during alcohol Pavlovian conditioning.**

**(A)** Mean ± standard error of the mean (SEM) time spent in the alcohol magazine during pre-CS periods across all experimental phases for photometry experiments. **(B)** Mean ± standard error of the mean (SEM) time spent in the alcohol magazine during pre-CS periods across conditioning sessions for optogenetics experiments. **(C)** Mean ± standard error of the mean (SEM) time spent in the alcohol magazine during pre-CS periods on the extinction session for optogenetics experiments.  Mean ± SEM alcohol leftover and g/kg of alcohol consumed during Conditioning sessions for photometry **(D)** and optogenetics **(E)** experiments.

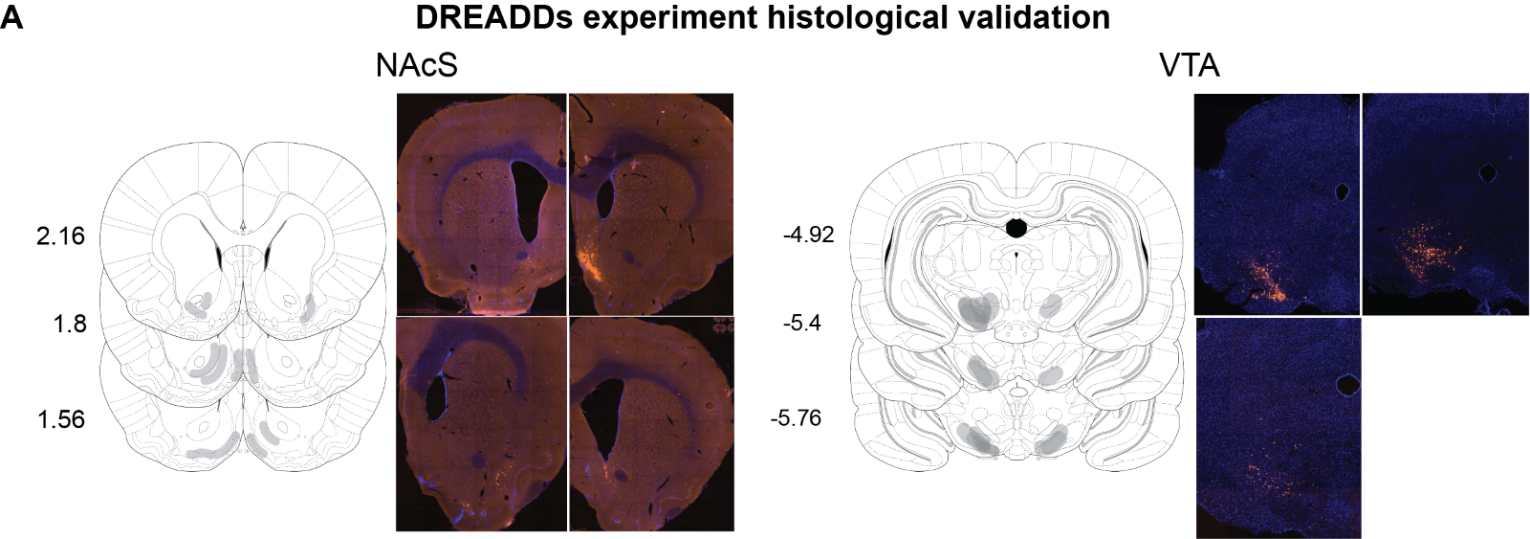

**Figure S3. Histological expression validation for DREADDs experiment.**

Expression validation for NAcS (left) and VTA (right) DREADDs injections, and representative images of sparse or off target expression.

 
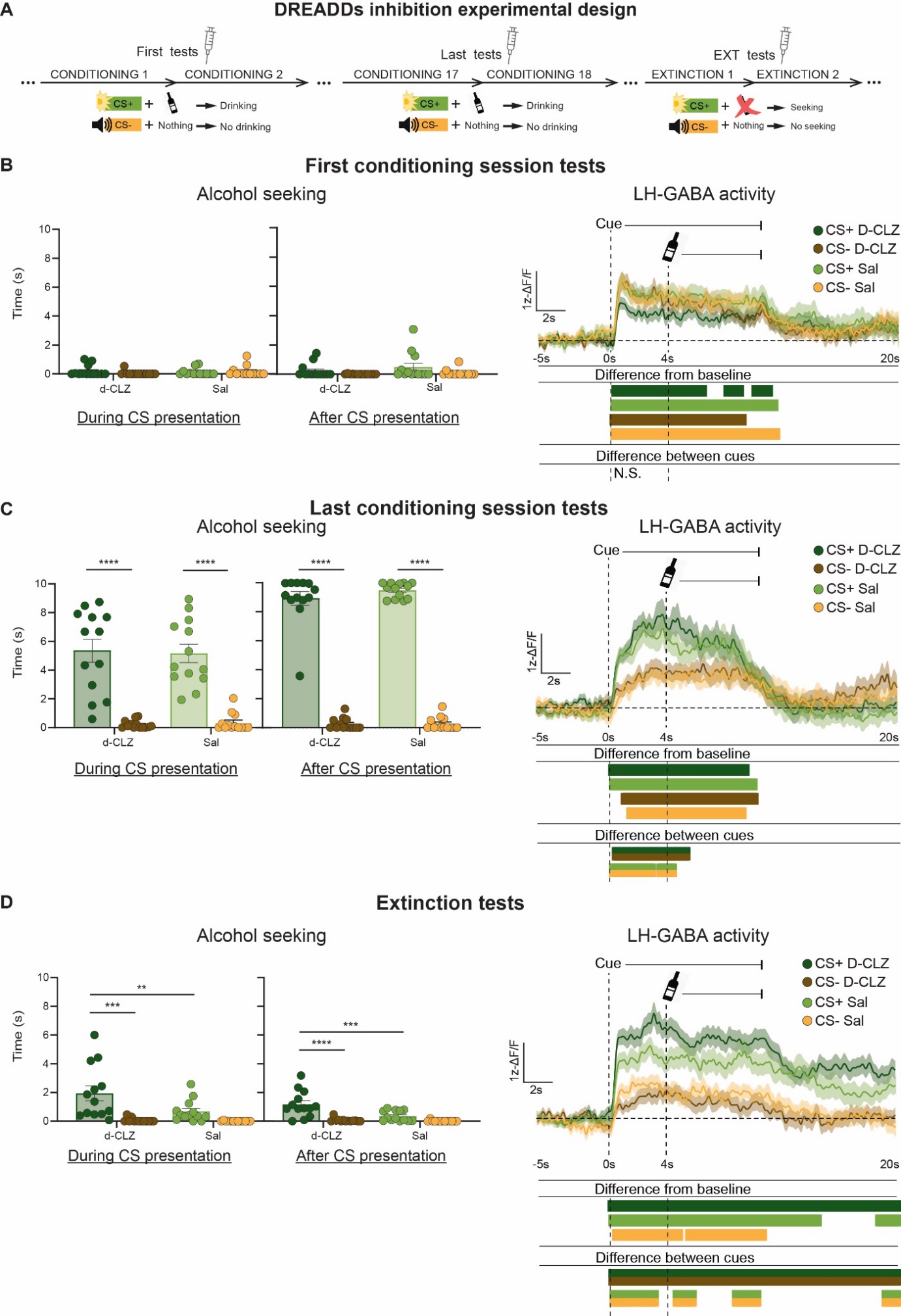

**Figure S4. Effect of sub-cutaneous deschloroclozapine injection on alcohol seeking, and LH-GABA activity.**

(**A**) Outline of the experimental procedure of the DREADDs test days, dots indicate normal training days (n = 13 females). **B-D** Mean ± standard error of the mean (SEM) time spent in the alcohol magazine during and after **(left)** CS presentation, and Ca2+ traces of LH-GABA activity **(right)** centered around cue onset (-5s to +20s) comparing LH-GABA activity to CS+ and CS- on the first conditioning **(B**; CS+ d-clz: n = 105, CS- d-clz: n = 105, CS+ sal: n = 106, CS- sal: n = 106**),** last conditioning (**C**; CS+ d-clz: n = 105 , CS- d-clz: n = 81 , CS+ sal: n = 105 , CS- sal: n = 81), and extinction **(D**; CS+ d-clz: n = 105, CS- d-clz: n = 80, CS+ sal: n = 106, CS- sal: n = 81**)** test days. For all photometry traces, bars at bottom of graph indicate significant deviations from baseline (dF/F ≠ 0), or significant differences between the specific events, determined via bootstrapped confidence intervals (99% CIs), and permutation tests with alpha 0.01 for comparisons between CS type and session. Vertical dashed lines indicate CS onset and alcohol delivery for CS+, horizontal line indicates baseline (dF/F = 0).

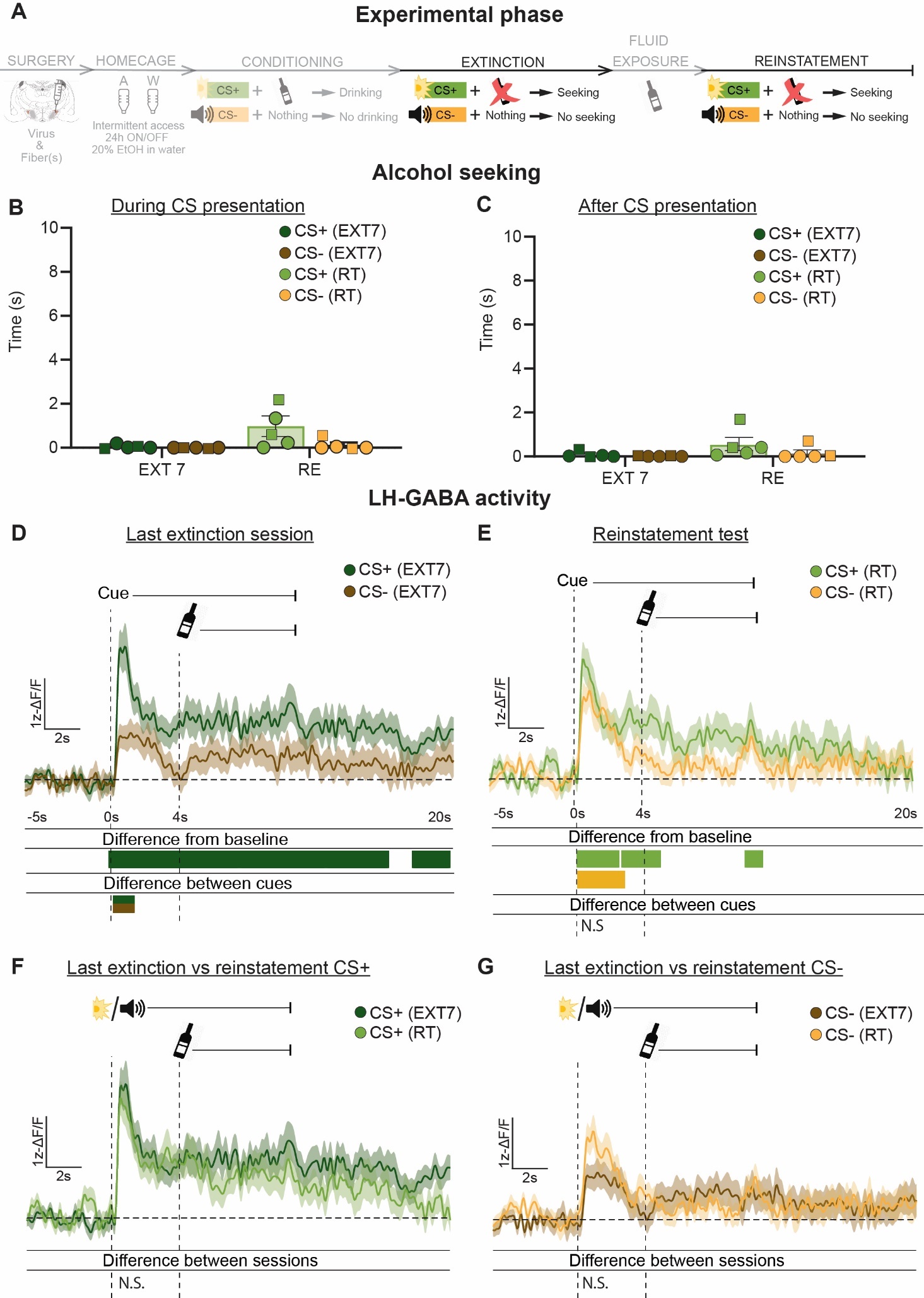

**Figure S5. Monitoring LH-GABA during non-primed reinstatement of alcohol seeking.**

(**A**) Outline of experimental procedure (n = 2 males, n = 3 females). Mean ± standard error of the mean (SEM) time spent in the alcohol magazine during (**B**) and after (**C**) CS presentation comparing the last extinction and reinstatement test sessions for the non-primed group. Ca2+ traces of LH-GABA activity centered around cue onset (-5s to +20s) comparing LH-GABA activity to CS+ and CS- on the last extinction (**D**) and reinstatement test (**E**) sessions. (CS+ EXT7: n = 56; CS- EXT7: n = 52; CS+ RT: n =56 ; CS- RT: n = 48). The same mean Ca2+ traces are plotted for comparisons across sessions between CS+ (**F**) and CS- (**G**).  For all photometry traces, bars at bottom of graph indicate significant deviations from baseline (dF/F ≠ 0), or significant differences between the specific events (CS+ EXT7; CS- EXT7; CS+ RT; CS- RT), determined via bootstrapped confidence intervals (99% CIs), and permutation tests with alpha 0.01 for comparisons between CS type and session. Vertical dashed lines indicate CS onset and alcohol delivery for CS+, horizontal line indicates baseline (dF/F = 0).

 
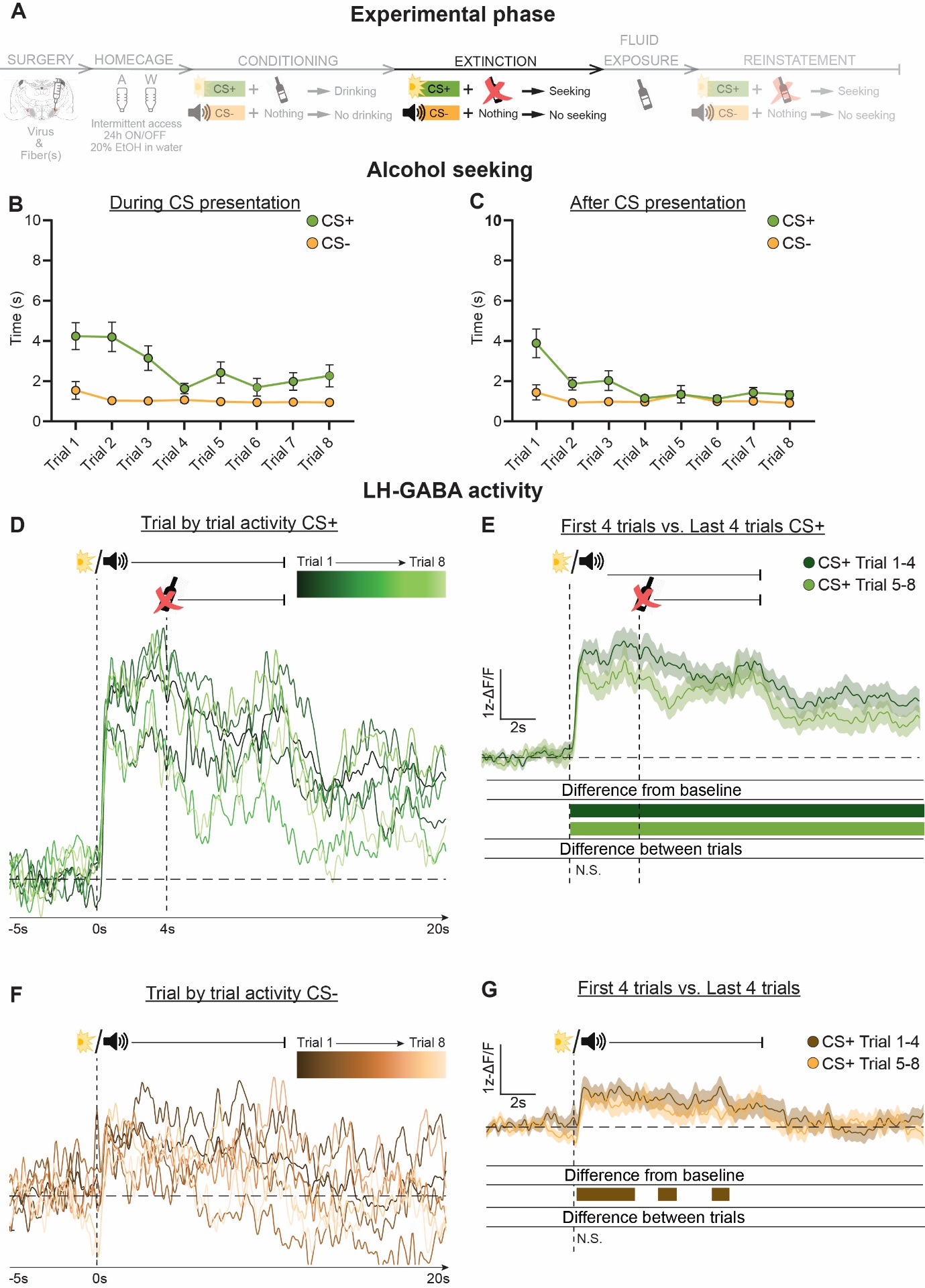

**Figure S6. Trial-by-trial alcohol seeking and LH-GABA activity during extinction.**

(**A**) Outline of experimental procedure (n = 2 males, n = 16 females). Mean individual trial time spent in the alcohol magazine during (**B**) and after (**C**) CS presentation during the first extinction session. Ca2+ traces of LH-GABA activity centered around cue onset (-5s to +20s) showing individual trial traces averaged across rats for CS+ (**D)** CS- (**F)**. Mean Ca2+ activity comparing the early (first 4) vs late (the last 4) trials for CS+ (**E)** and CS- **(F)** on the first Extinction session. For all average traces comparing early vs late activity, bars at bottom of graph indicate significant deviations from baseline (dF/F ≠ 0), or significant differences between the specific events (CS+ Trial 1-4 vs. CS+ Trial 5-8; CS- Trial 1-4 vs. CS- Trial 5-8), determined via bootstrapped confidence intervals (99% CIs), and permutation tests with alpha 0.01 for comparisons between early and late activity. Vertical dashed lines indicate CS onset and alcohol delivery for CS+, horizontal line indicates baseline (dF/F = 0).

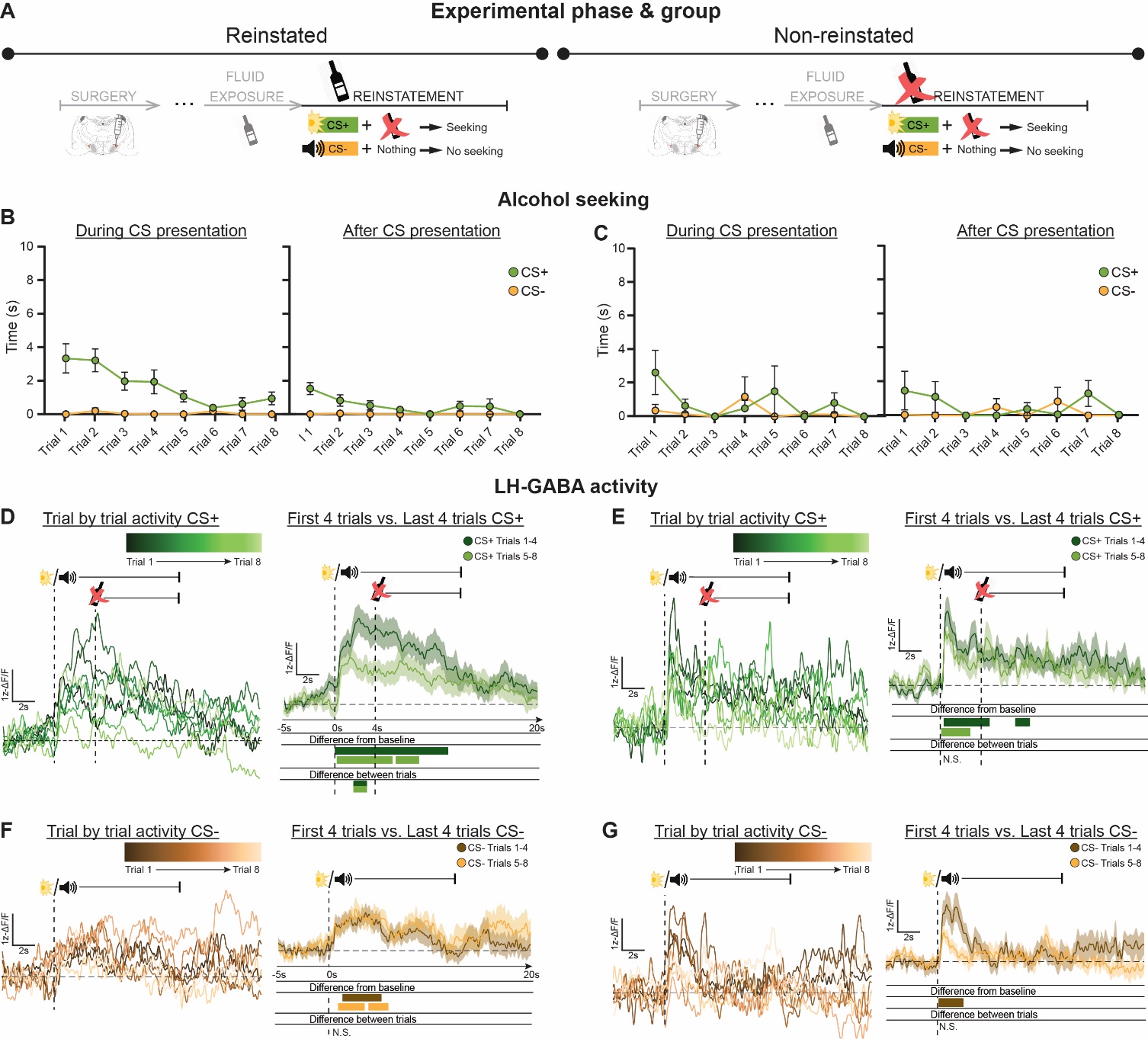
**Figure S7. Trial-by-trial alcohol seeking and LH-GABA activity during reinstatement.**

(**A**) Outline of experimental procedure (n = 2 males, n = 16 females). Trial-by-trial alcohol seeking and Ca2+ activity, and early (first 4) vs late (the last 4) trial comparisons for the reinstated (Left plots) and non-reinstated (Right plots) groups. Mean individual trial time spent in the alcohol magazine during (**B,C, left**) and after (**B,C, right**) CS presentation during the reinstatement sessions for the reinstated **(B)** and non-reinstated **(C)** groups. Ca2+ traces of LH-GABA activity centered around cue onset (-5s to +20s) showing individual trial traces averaged across rats for CS+ (**D, E, left)** CS- (**F, G, right)**. Mean Ca2+ activity comparing the early (first 4) vs late (the last 4) trials for CS+ (**D, E, right)** and CS- **(F, G, left)** on the first Extinction session. For all average traces comparing early vs late activity, bars at bottom of graph indicate significant deviations from baseline (dF/F ≠ 0), or significant differences between the specific events (CS+ Trial 1-4 vs. CS+ Trial 5-8; CS- Trial 1-4 vs. CS- Trial 5-8), determined via bootstrapped confidence intervals (99% CIs), and permutation tests with alpha 0.01 for comparisons between early and late activity. Vertical dashed lines indicate CS onset and alcohol delivery for CS+, horizontal line indicates baseline (dF/F = 0).

**Supplementary Table 1.** *Time periods (in seconds) where the photometry statistical tests are significant. Time 0 refers to the start of the 10 second Cue Period.*

| Experimental Phase | Figure | Statistical Comparison | Significance time window  (Seconds; 0 = Cues turned on) |
| --- | --- | --- | --- |
| Conditioning | 3D | Bootstrapped CI, First Cond. session CS+ | 0.31🡪11.76 |
|  | 3D | Bootstrapped CI, First Cond. session CS- | 0.31🡪10.94 |
|  | 3D | Permutation tests: First Cond. session (CS+ v CS-) | n.s. |
|  | 3E | Bootstrapped CI: Last Cond. session CS+ | 0.31🡪9.62 |
|  | 3E | Bootstrapped CI: Last Cond. session CS- | 0.44🡪10.6 |
|  | 3E | Permutation tests: Last Cond. session (CS+ v CS-) | 0.44🡪6.9 |
|  | 3F | Permutation tests: CS+ (First v Last Cond. session) | 2.14🡪7 |
|  | 3G | Permutation tests: CS- (First v Last Cond. session) | 0.5🡪1.57 |
| Extinction | 4E | Bootstrapped CI, First Ext session CS+ | 0.25🡪20 |
|  | 4E | Bootstrapped CI, First Ext session CS- | 0.44🡪8.6; 10🡪10.12; 10.37🡪10.81 |
|  | 4E | Permutation tests: First Ext. (CS+ v CS-) | 0.31🡪20 |
|  | 4F | Permutation tests: CS+ (Last Cond. v First Ext.) | 9.4🡪12.9; 16.8🡪18.42 |
|  | 4G | Permutation tests: CS- (Last Cond. v First Ext.) | n.s. |
|  | 5E | Bootstrapped CI, Last Ext. session CS+ | 0.25🡪20 |
|  | 5E | Bootstrapped CI, Last Ext. session CS- | 0.44🡪2.57; 8.5🡪8.9 |
|  | 5E | Permutation tests: Last Ext. session (CS+ v CS-) | 0.44🡪1.64; 2.58🡪5.03 |
|  | 5F | Permutation tests: CS+ (First Ext. v Last Ext.) | 1.13🡪5.47; 7.23🡪11.5 |
|  | 5G | Permutation tests: CS- (First Ext. v Last Ext.) | n.s. |
| Reinstatement | 6D | Bootstrapped CI, Last Ext. CS+ | 0.38🡪9.24 |
|  | 6D | Bootstrapped CI, Last Ext. CS- | n.s. |
|  | 6D | Permutation tests: Last Ext. session (CS+ v CS-) | 2.64🡪4.91 |
|  | 6E | Bootstrapped CI, Reinstatement CS+ | 0.38🡪9.87 |
|  | 6E | Bootstrapped CI, Reinstatement CS- | 0.57🡪6.10; 7.48🡪9.87 |
|  | 6E | Permutation tests: Reinstatement (CS+ v CS-) | 5.59🡪7.67 |
|  | 6F | Permutation tests: CS+ (Last Ext v Reinstatement) | n.s. |
|  | 6G | Permutation tests: CS- (Last Ext v Reinstatement) | 2.58🡪3.33 |
|  | S7D | Permutation tests: CS+ (Reinstatement Trials 1-4 v 5-6) | 2.14🡪3.4 |
